# Global patterns of natural selection inferred using ancient DNA

**DOI:** 10.64898/2026.01.07.697984

**Authors:** Laura L. Colbran, Jonathan Terhorst, Iain Mathieson

## Abstract

Ancient DNA has revolutionized our understanding of human history, and is now yielding important insights into evolution and natural selection. However, studies of selection using ancient DNA have largely been limited to Europe, excluding populations in other parts of the world. While many selective pressures were local to specific populations others, for example those related to the development of agriculture, may have been universal. By studying a broader range of global populations, we can identify examples of local adaptation but also more general principles of adaptation to climatic, social and technological changes.

We leveraged ancient DNA to test for selection in 7244 individuals from 13 ancient and 19 present-day populations across five regions: Europe, East Asia, South Asia, Africa and the Americas. In each region, we tested for selection using multiple approaches that account for complex demographic histories. We identify 34 genome-wide significant signals of selection, including both known and novel loci. We find a high degree of shared signal across regions, suggesting extensive parallel or shared adaptation. Using a novel admixture-aware time series method, we find that the strength of selection on many variants varied with time and ancestry, for example increasing selection at *ADH1B* in East Asia over the past few thousand years, and differential selective pressures across Europe at *HERC2*.

Finally, we developed a test for polygenic selection on complex traits by modeling the frequencies of trait-associated alleles identified in GWAS. We tested for selection jointly across regions, avoiding the confounding effect of population stratification by excluding the European or East Asian GWAS population from the selection test. We find evidence for directional selection on immune traits, highlighting the importance of infectious disease in recent human evolution.

## Introduction

As modern humans migrated out of Africa and throughout the world over the last 50,000 years, they encountered dramatically different environmental pressures, providing the opportunity for genetic adaptation to shape human variation. In the past 10,000 years (the Holocene), populations in different parts of the world independently developed agriculture, urbanism and new forms of social organization. This raises the question of how humans adapted to these new environments, how much those adaptations were shared across populations, and how they affect human diversity today. Understanding which parts of the genome have been subject to selection can highlight functionally important regions, and contribute to understanding how they influence phenotype. Further, strongly selected regions can play a role in medically-significant phenotypes, regardless of their current fitness cost, making patterns of selection informative for understanding variation in health outcomes among populations.

The emerging science of ancient DNA has brought a new perspective to studies of human evolution. The ability to observe allele frequency changes directly combined with a better understanding of demographic history has enabled powerful inference of natural selection [1], precise estimates of timing [2, 3], and the ability to recover signatures of selection that have been obscured by admixture or genetic drift [4, 5]. However, due to sampling limitations, large-scale ancient DNA selection studies have been limited to Europe [1, 3, 6–8], limiting the scope for comparative evolutionary studies.

We therefore took advantage of the recent influx of non-European ancient DNA to conduct a scan for selection across five regions of the world—Africa, East Asia, Europe, South Asia, and the Americas— each with hundreds of ancient samples and present-day population data. Each of these regions has a complex history of Holocene admixture and we adapted an allele frequency-based test [1] to account for more general admixture history. We further adapted this test to detect polygenic selection, by modeling expected frequencies of alleles associated with a complex trait. One of the primary advantages of ancient DNA is that ancient individuals can be directly dated so we also developed a time series approach to model the evolution of allele frequencies in admixed populations. We find evidence of both private and shared targets of selection, demonstrating the power of ancient DNA to highlight the interconnected nature of evolution in the recent human past.

## Results

### An allele frequency-based test for selection

Identifying natural selection in genomic data requires distinguishing its effects from the effects of admixture and genetic drift. The Holocene is characterized by high levels of admixture between previously diverged populations, which can lead to both false positives and false negatives in selection scans [4, 5, 8]. We therefore developed a test where we model expected allele frequency based on genome-wide admixture proportions, extending a previous approach [1] by modeling the ancient populations themselves as admixtures of unobserved source populations. This approach is similar to that of Cheng et al. [9], except that we treat source populations as independent. We perform a likelihood ratio test of whether each SNP is consistent with the fitted admixture model and then apply a correction for the inflation in test statistics due to genetic drift by using genomic control. This test detects deviations in allele frequency due to selection in the ancestors of one or more of the sampled ancient or modern populations. If the admixture model is poorly specified, the test might misattribute selection that occurred in the ancestors of the source populations to the test populations, but should not identify selection where none occurred. We applied this method to test for selection at 1,150,640 variants captured by the 1240k reagent [10, 11] for five regions with a genetic record which includes both ancient and present-day samples: Africa, East Asia, Europe, South Asia, and Central/South America (Fig. 1A). In each region, we defined populations of ancient and present-day individuals based on genome-wide ancestry and inferred ancestry proportions from the unobserved source populations using ADMIXTURE [12] (Figures S1-S5).

**Figure 1.**
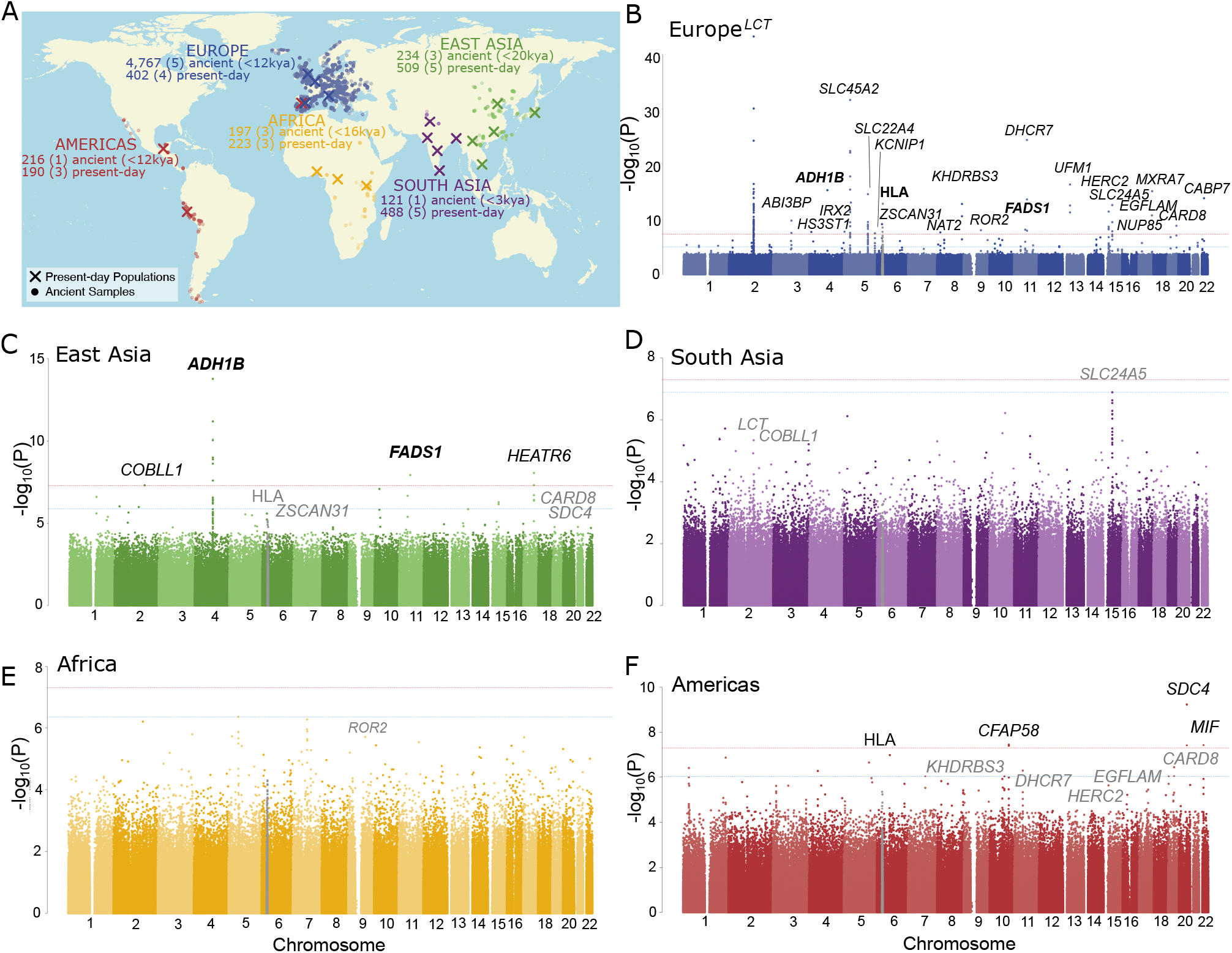
Scan for selection on 1240k variants based on admixture proportions A) for 5 regions with ancient individuals (circular points) and present-day populations (X = sampling locations). Manhattan plots of the results for B) Europe, C) East Asia, D) South Asia, E) Africa, and F) the Americas. Significant SNPs are filtered to only include those with at least one more nearby (at least 2 more for Europe), in high LD, and also passing FDR*<*0.1 (Methods). The HLA is indicated in grey, as it was treated independently of the rest of the genome. Peaks are labeled with the causal gene if known, or the nearest to the lead SNP. Bold text indicates loci passing genome-wide multiple testing in multiple regions, while grey text labels loci that are genome-wide significant in another region, and nominally significant in the indicated region after correcting for the total number of genome-wide significant peaks.

### 34 signals of selection with extensive sharing across continents

After filtering out SNPs with potential genotype errors, and performing a separate multiple testing correction at the HLA (Figures S6 and S7), we identified 25, 4, and 5 genome-wide significant (*p <* 5 × 10^*−*8^) peaks in Europe, East Asia, and the Americas respectively, with none in South Asia or Africa (Fig. 1B-F, Table S1, Figure S8). The differing numbers of peaks reflect the relative power we have in each region; simulations show that we have the most power to detect selection in Europe–though still only around 50% for a selection coefficient of 0.02—and the least in Africa (Supplementary Note A). Comparison with other selection scans conducted using European data suggests that our ordering of loci is similar, and the main factor affecting the number of loci reported is the difference in the genome-wide significance threshold (Supplementary Note B).

The loci we detect include well-known signals of selection associated with diet, pigmentation and the immune system, as well as novel loci of unknown function (Table S1, Supplementary Note C). Three loci were genome-wide significant in more than one region (*ADH1B* and *FADS1* in both Europe and East Asia, and HLA-B in Europe and the Americas), so our 34 signals correspond to 31 distinct loci. Additionally, thirteen were genome-wide significant in one region and nominally significant (at P*<*0.05/31) in at least one other. Variants passing FDR*<* 0.05 (*N* = 215) in Europe, where we had the highest power, are significantly enriched in the lower tail of the p-value distribution for all other non-African regions (P = 9.0 × 10^*−*8^, 1.5 × 10^*−*5^, 9.2 × 10^*−*11^ and 0.45 for East Asia, the Americas, South Asia, and Africa, respectively; Kolmogorov-Smirnov test). Overall, this suggests that a relatively high proportion of selected variants have shared signals across regions, reflecting either parallel selection on standing variation or selection on distinct variants at the same loci. One caveat is that our admixture model is only an approximation to the true demographic history, and selection in a common ancestral population might be incorrectly attributed to multiple test populations.

Closer inspection of the signals can resolve these possibilities. For example, at *ADH1B* —an alcohol dehydrogenase gene where a variant associated with poor alcohol metabolism is known to have been under strong selection in East Asia [13–16]—the putatively causal East Asian variant (rs1229984) is rare in Europe, indicating selection on distinct variants. Similarly, the signal of selection in the Americas— replicating in East Asia—in the region around *SDC4* overlaps eQTLs for multiple immune-associated genes, including members of the *WFDC* gene cluster. In contrast, other shared loci including *FADS1*, where a gene regulatory change affecting fatty acid metabolism was selected in both East Asia and in Europe, and *SLC24A5* affecting skin pigmentation in Europe and South Asia (Figure S9), involve selection acting on the same haplotype.

### Admixture-aware modeling of time series data

To be able to further disentangle these selection signals, we extended our previous time-series modeling approach [`bmws;` 2] to take into account varying levels of admixture among individuals. We model allele frequencies evolving separately in each admixture component, allowing selection coefficients to vary over time and across admixture components, while controlling for changes in allele frequency due to admixture. We conducted simulations to validate the approach (Supplementary Note D) then applied it to all significant loci. This allows us to identify changes in the strength of selection over time. For example, at *ADH1B* we find a distinct increase in the selection coefficient between 100 and 150 generations ago (Fig. 2A-C).

**Figure 2.**
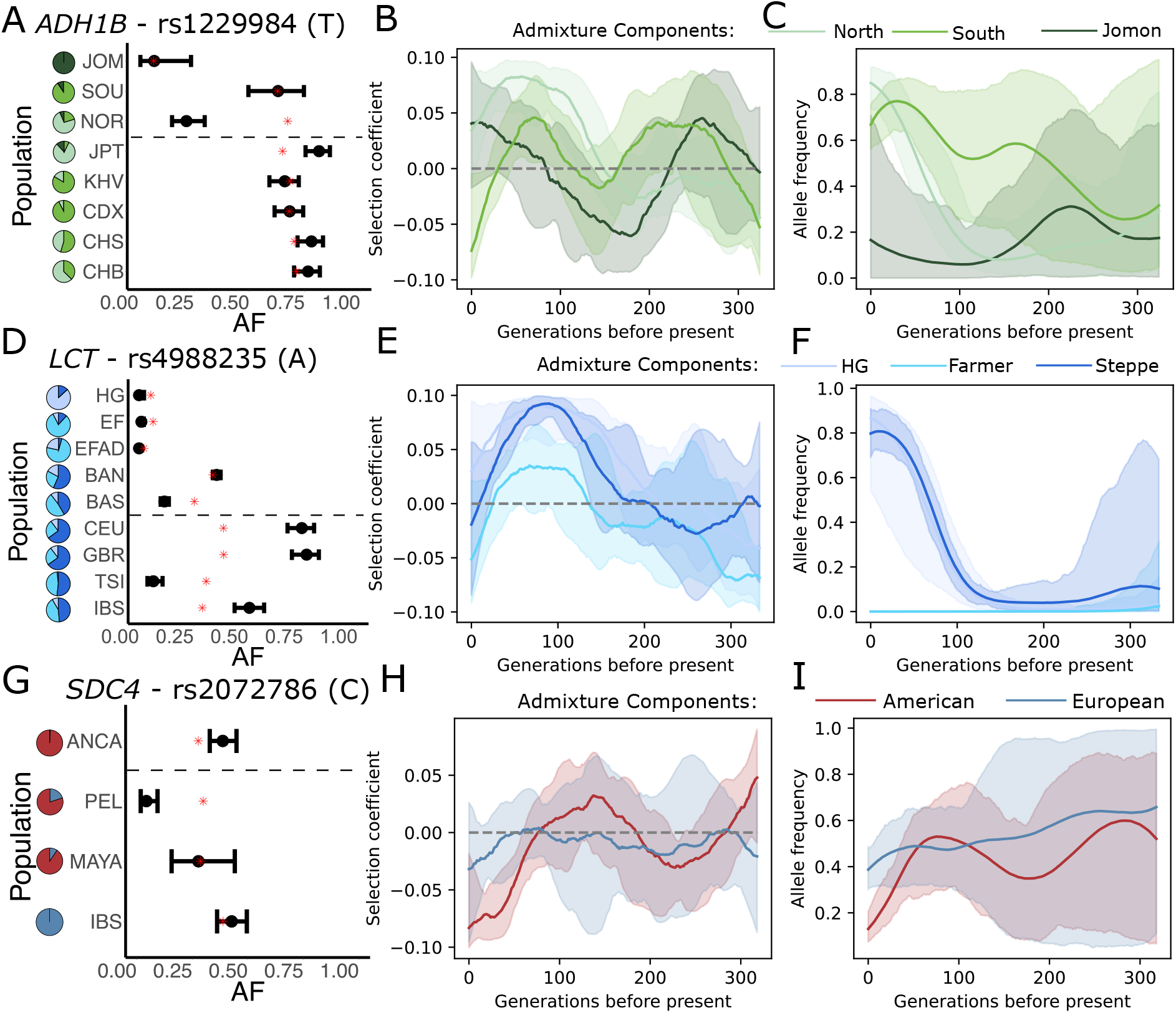
Ancestry-specific time series of the most strongly selected loci in East Asia, Europe and the Americas. A) Observed allele frequencies; B) inferred selection coefficients; C) Allele frequency trajectories in admixture components at *ADH1B* in East Asia, D-F) *LCT* in Europe, and G-I) *SDC4* in the Americas. We plotted the information for the derived allele of the lead SNP in each peak. In the frequency plots (first column), the bars represent 95% confidence intervals, while the red asterisk indicates the fitted frequency from the admixture model and the dashed line separates ancient (top) from present-day (bottom) groups. The pie charts indicate the average proportion of each admixture component for each population. See Methods for population abbreviations. In the plots of inferred selection coefficient and allele frequency (second and third columns), different colors distinguish the *k* source populations, solid lines represent fitted values, and shaded areas represent the 5-95% posterior credible intervals.

At rs4988235 in *LCT*, the strongly selected variant associated with adult lactase persistence in Europe [17], we infer strong selection up to about 100 generations ago, consistent with previous estimates of timing [2, 3] (Fig. 2D-F). Differences in selection coefficients across ancestry components likely reflect differences across geographic or social structure that is correlated with ancestral structure. For example, we also find smaller estimates of s in the Farmer ancestry component suggesting little selection on lactase persistence in populations with high Farmer ancestry (e.g. TSI and other southern Europeans). Even in Northern Europe, selection drops off into the present-day, consistent with several lines of evidence showing little or no recent selection [18, 19]. In contrast, at the lead SNP at *SDC4* in the Americas, the time series suggests relatively little selection until a spike in selection against the derived allele in the American component in the last 80 generations, reflected by the lower-than-expected frequency in present-day Peruvians (Fig. 2G-I).

### Parallel adaptation of fatty acid metabolism in Europe and East Asia

Selection for the same *FADS1* haplotype has been previously identified in both Europe [1] and East Asia [20, 21]. The selected haplotype increases the rate of synthesis of plasma unsaturated fatty acids, is believed to be an adaptation to a relatively plant-heavy agricultural diet [22], and appears to still be under selection today in Europe [23]. We estimate that selection in Europe remained relatively constant through time in the Farmer ancestry component with a selection coefficient around 1% (Fig. 3A-C). In contrast, in East Asia, we find that selection has intensified over time, with the allele frequency increasing substantially in the past 100 generations (Fig. 3D-F).

**Figure 3.**
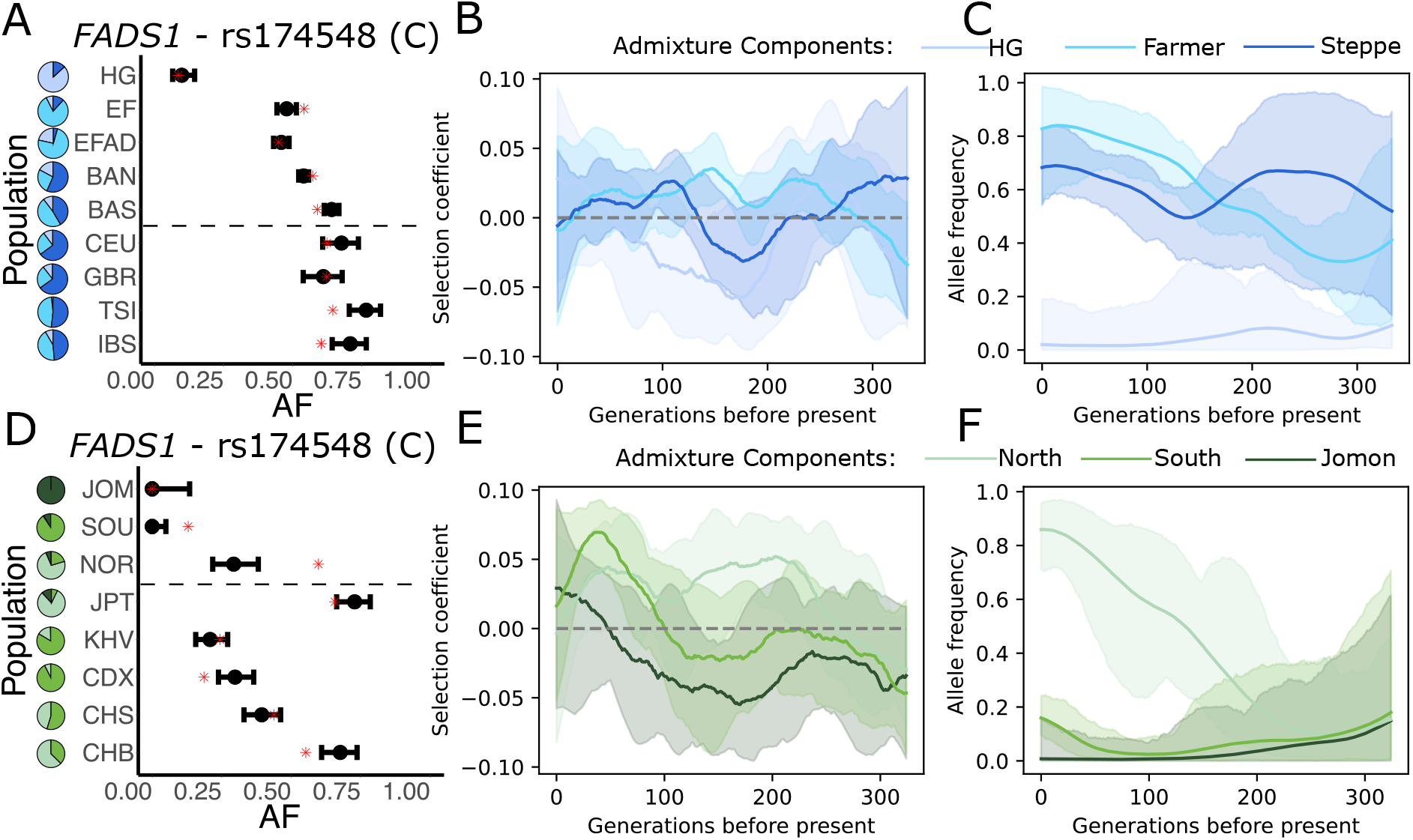
Ancestry-specific time series of *FADS1* in Europe and East Asia. A) Observed allele frequencies; B) inferred selection coefficients; C) Allele frequency trajectories in admixture components at *FADS1* in Europe and D-F) in East Asia. Details of the plots are the same as in Fig. 2

### Selection on pigmentation in Europe

The SNP rs12913832 in *HERC2* is the major determinant of blue (versus brown) eye color through its regulatory effects on *OCA2*, and is a known target of selection in Europe [1, 24]. We detect both divergent selection and a persistent difference in frequency across ancestry groups (Fig. 4A-C), indicating long-term geographically structured balancing selection consistent with the North-South frequency gradient that has persisted from the Mesolithic to the present-day [25]. It appears therefore that light eye color was beneficial in Northern Europe and deleterious in southern Europe, and that this selection gradient persisted through at least two almost total population replacements. In contrast, at skin pigmentation loci *SLC45A2* and *SLC24A5*, we find consistent selection driving allele frequencies almost to fixation on all ancestry backgrounds (Fig. 4D-I). Selection on skin pigmentation is thought to reflect pressure for increased vitamin D synthesis [26]. We find that selective pressure on *DHCR7* [27], which encodes a key part of the vitamin D synthesis pathway, intensified in the past 100 generations (Fig. 4J-L), consistent with our previous findings from Britain [2] that selective pressure on vitamin D levels persisted or even intensified into the past few thousand years.

**Figure 4.**
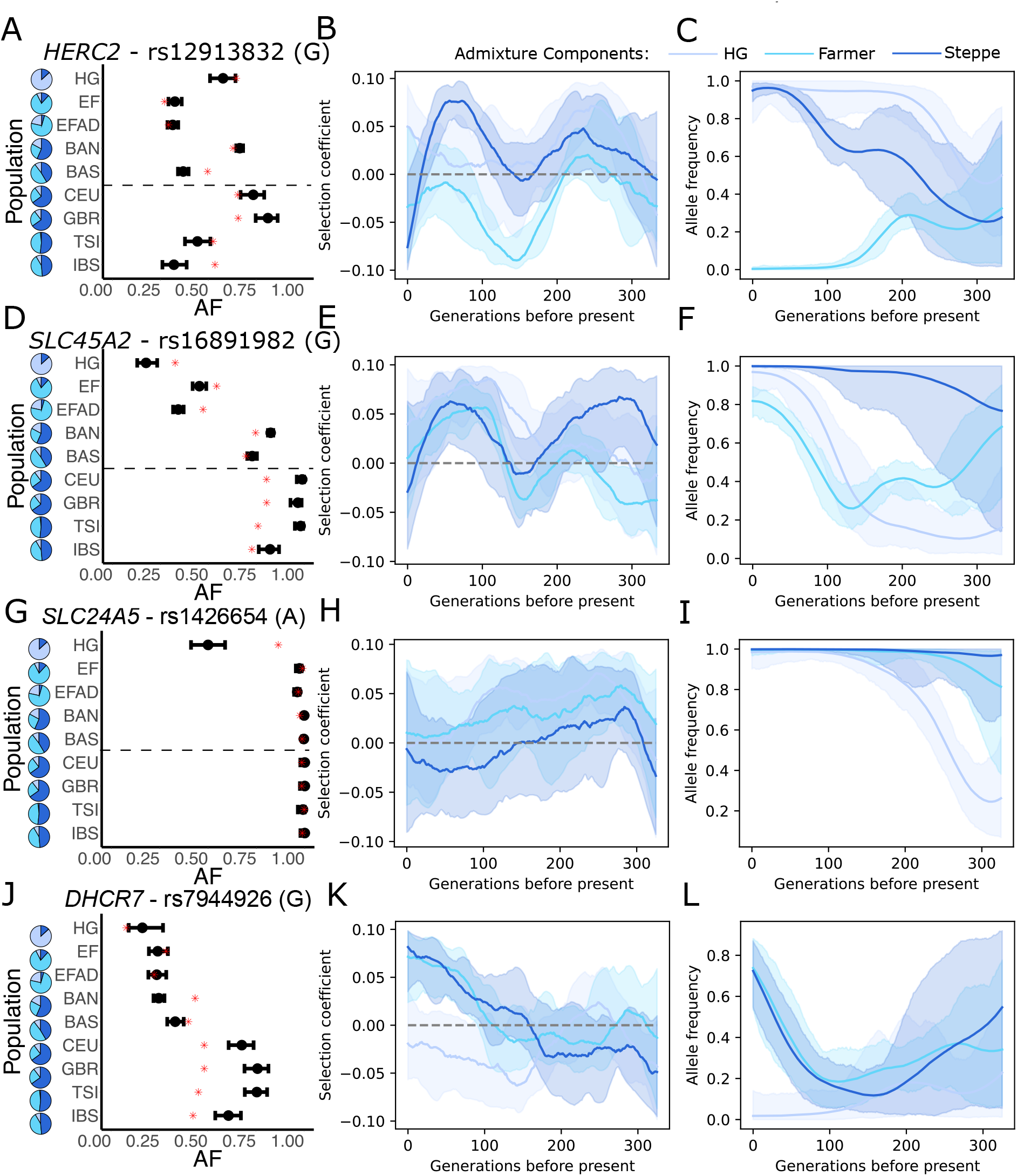
Details of the *HERC2, SLC45A2, SLC24A5* and *DHCR7* loci in Europe. A) Plot of observed Allele Frequencies, B) Inferred Selection coefficients, and C) AF trajectories in admixture components at *HERC2*, followed by D-F) *SLC45A2*, G-I) *SLC24A5*, and J-L) *DHCR7*. We plotted the information for the derived allele of the lead SNP in each peak. Details of the plots are the same as in Fig. 2.

### Widespread directional selection on immunity

Most human complex traits are highly polygenic. To study these traits, we extended our allele frequency-based test to detect polygenic selection. We first identify trait-associated alleles based on summary statistics from genome-wide association studies (GWAS). For each ancient and modern population, we then count the proportion of trait increasing alleles, across all variants and individuals in each population. We treat this proportion as though it were an allele frequency, and perform the same likelihood ratio test, conditional on admixture proportions, that we used for the per-SNP analysis.

There are two advantages to this approach. First, since it only counts observed variants, it is insensitive to genotypes missing at random. Second, we can compute a well-calibrated P-value by permuting the effect directions of variants. Small P-values indicate that the distribution of trait-associated variants is very different than expected based on genome-wide admixture proportions indicating directional selection in one or more populations. Conversely, large P-values indicate that the frequencies of trait-associated variants are “too close” to the genome-wide admixture proportions, indicating that the trait is under stabilizing selection (Figure S10). However, this can only happen when the tested variants capture a large fraction of the heritability of the trait, which is not the case here [28] (Figure S11).

A persistent problem with tests of polygenic selection is that they rely on variants identified in GWAS which can be confounded by population stratification, leading to false-positive signals of selection [29, 30]. One solution to this problem is to only perform selection tests on summary statistics obtained in a different population from the one on which the selection test is performed [31]. We therefore used summary statistics obtained in European populations to test for selection in all populations except Europe (Fig. 5).

**Figure 5.**
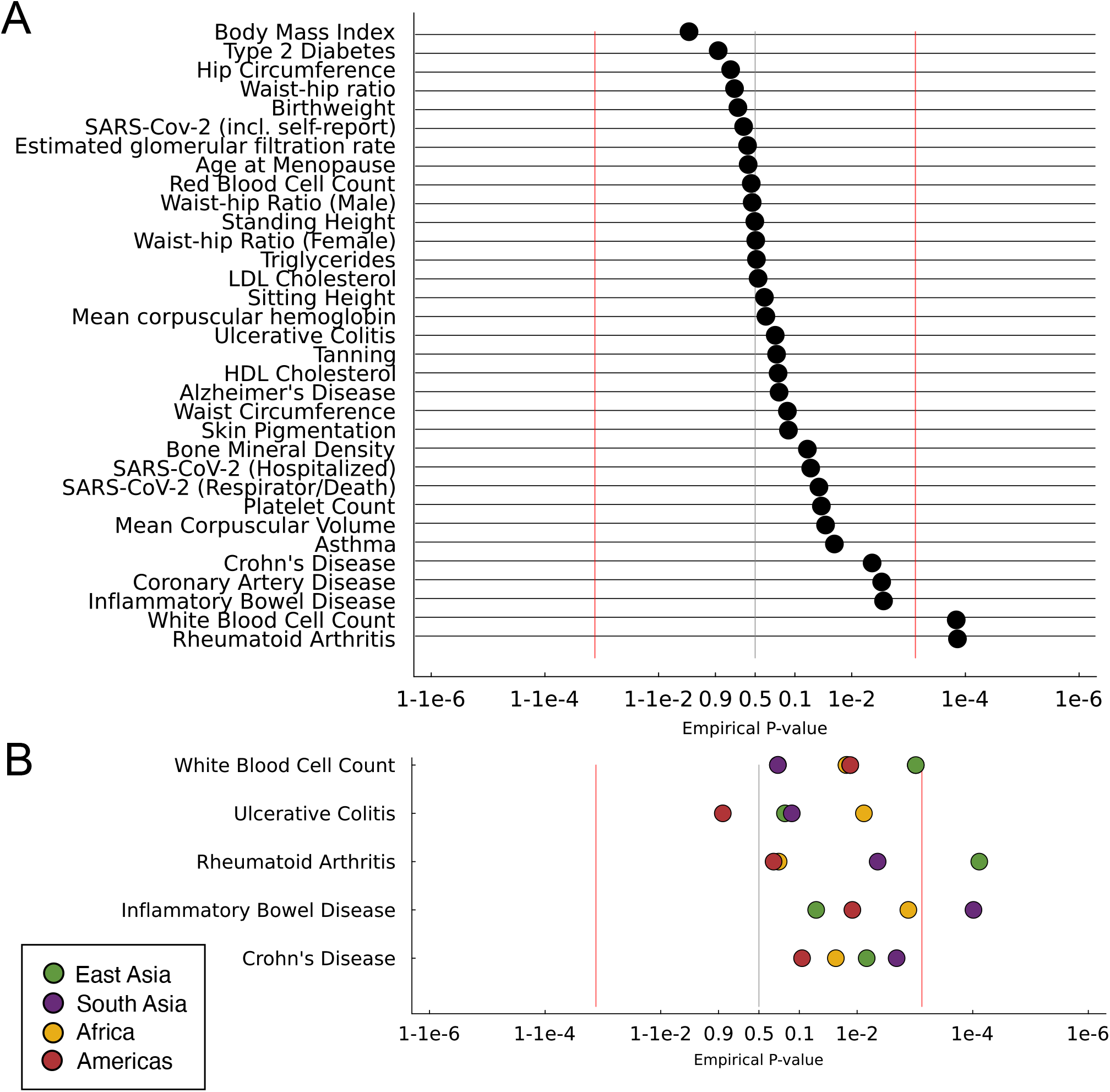
Several traits show evidence for polygenic selection. A) We tested for polygenic selection across SNPs from European-ancestry GWAS for 33 traits in jointly across 4 regions, excluding Europe to avoid the possibility of population stratification inducing false positives. B) Region-specific results for immune traits with significant evidence for selection. The X-axis is plotted on a log scale, as −*log*_10_(*p*) + *log*_10_(0.5) where *p <* 0.5 and log_10_(1*− p*) *−* log_10_(0.5) for *p >* 0.5. Red lines indicate a 2-sided Bonferroni correction threshold.

Testing 33 complex traits, we find a highly significant signal of directional selection on rheumatoid arthritis (P = 1.4 × 10^*−*4^) and white blood cell count (P = 1.5 × 10^*−*4^). If we include Europe in the analysis, (Figure S12) we additionally find directional selection on skin pigmentation, mean corpuscular hemoglobin (MCH) and Crohn’s diease. Although these are potentially affected by stratification, they do appear to reflect the effect of directional selection, as shown by other lines of evidence for selection on skin pigmentation and immune traits.

## Discussion

Our results highlight several key aspects of recent natural selection in humans. First, we confirm that classic (i.e. strong, complete) selective sweeps are rare in recent human evolution [32]. Second, for many of the partial selective sweeps that do exist, the strength of selection varies with time, space and ancestry. Third, many signals of selection are shared between populations. In some cases (e.g. *FADS1*) the same haplotype was selected in different populations, while in others (e.g. *ADH1B, SDC4*), different haplotypes at the same locus were selected. Similarly, the strongest signals of polygenic selection we detected—of directional selection on immunity—are largely shared across populations.

These shared signals of selection may reflect adaptation to shared selective pressures, for example the adoption and development of agriculture independently in many parts of the world. For example, the strongest signals of selection at individual loci (*LCT, ADH1B, FADS1*) are all plausibly related to the consumption of agricultural products. Shared signals may also reflect the fact that some loci might be repeatedly targeted by sweeps. For example, genes involved in immunity might experience repeated sweeps, or parallel sweeps in different populations, even if different pathogens are involved. Finally, some shared signals might reflect imperfect modeling; one limitation of our approach is that it does not distinguish between selection in the history of ancient or present-day populations and selection acting in ancestral populations that is not fully captured by the inferred components. In particular because the components are inferred, and only approximately capture the relevant demography, selection acting in deeper ancestral groups may not be fully captured by the model, leading to residual signals of selection that might be shared across groups.

Skin pigmentation in Europe is well-known to have been under selection, probably in response to differing UV exposure at higher latitudes and the downstream effects on Vitamin D biosynthesis [26, 33]. We confirm this result, but we do not see a clear signal in other populations. For example, in East Asia, none of the genome-wide significant loci are clearly related to pigmentation and we see no polygenic signal of selection on skin pigmentation using either European or East Asian summary statistics. We do see a suggestive signal of polygenic selection on tanning response, raising the hypothesis that adaptation to latitude in East Asia focused on plastic response rather than constitutive skin pigmentation as in Europe.

The immune system is another key target of selection. This is reflected by multiple signals at the HLA, and significant directional selection on loci influencing Rheumatoid Arthritis, Crohn’s disease and other immune traits. RA is an autoimmune disease with a large role played by inflammatory pathway including HLA genes [34]. While RA itself is likely not the source of selective pressure, it is an example of a clinical trait being influenced by evolution; pro-inflammatory signals may be beneficial in the case of an infection, but cause problems in other contexts.

The main limitation of our study is that we are still relatively underpowered, particularly outside Europe. Even in Europe, power drops off rapidly for selection coefficients between 1-2%. While the number of loci we detect is broadly consistent with other approaches using comparable significance thresholds [1, 3, 5–7], it seems likely that there were many more variants under selection, with selection coefficients less than 1%. Indeed, some other scans (e.g. [8]) find exactly that (Supplementary Note B). However, the questions of how accurately those sweeps can be identified, how much they are shared across populations, and how much they target *de novo* or standing variation remain unresolved.

## STAR Methods

### METHOD DETAILS

#### Population samples and admixture models

We tested for selection in 5 regions that had genetic representation of both ancient and present-day populations: Africa, East Asia, South Asia, Europe and Central/South America (Fig. 1A). Most data were obtained from the Allen Ancient DNA Resource (AADR) [35], with additional samples from the HGDP [36] where necessary. For each region, we projected samples into PCA space constructed using samples genotyped on the Human Origins array from the same regions, then identified groups of genetically-similar ancient samples that encompassed the present-day populations. We defined those groups by drawing polygons around them in PCA space. As most ancient samples were pseudohaploid, we used pseudohaploid versions of samples from the 1000 Genomes project, reported in the AADR.

In Africa, we included the Yoruba (YRI; *N* = 101) and Luhya (LWK; *N* = 101) populations from the 1000 Genomes Project (1kG) [37], as well as Biaka from the Human Genome Diversity Project (HGDP;BIAK; *N* = 21) [36]. We identified 3 distinct ancient groups corresponding to Central Africa (CENT; *N* = 33), East Africa (EAST; *N* = 149), and those genetically similar to present-day Bantu speakers (BANT; *N* = 15) (Figure S1). We used *K* = 3 for ADMIXTURE modeling. The ancient samples used here were originally published in [38–45]. To construct the PCA we used Human Origins samples published in [44, 46–50].

For East Asia, we represented present-day populations with five from 1kG: Han Chinese from Beijing (CHB; *N* = 103) and Southern China (CHS; *N* = 106), Chinese Dai (CDX; *N* = 99), Japanese (JPT; *N* = 104), and Vietnamese (KHV; *N* = 97). We had 3 ancient groups corresponding to Northern China (NOR; *N* = 103), Southern China (SOU; *N* = 107), and the Jomon in Japan (JOM; *N* = 24), and used *K* = 3 for ADMIXTURE (Figure S2). The ancient samples used here were originally published in [51–64]. To construct the PCA we used Human Origins samples published in [46, 48, 65, 63].

For South Asia we used Gujarati (GIH; *N* = 105), Punjabi (PJL; *N* = 96), Telugu (ITU; *N* = 103), Tamil (STU; *N* = 99), and Bengali (BEB; *N* = 85) populations from 1kG, and kept the ancient samples in 1 group (ANCS; N =121). We used k=2 for ADMIXTURE (Figure S3). The ancient samples used here were originally published in [66, 67]. To construct the PCA we used Human Origins samples published in [46, 48, 63, 68].

For present-day European populations we used Italian Toscani (TSI; *N* = 108), Iberian (IBS; *N* = 103), British (GBR; *N* = 92) and Northwestern Europeans (CEU; *N* = 99) from 1kG, and split the ancient samples into 5 groups. These were hunter-gatherers (HG; *N* = 157), early farmers (EF; *N* = 605), admixed early farmers (EFAD; *N* = 892), northern Bronze Age (BAN; *N* = 2113), and southern Bronze Age (BAS; *N* = 1000). We used k=3 for ADMIXTURE (Figure S4). The ancient samples used here were originally published in [1, 11, 19, 25, 52, 67, 69–147]. To construct the PCA we used Human Origins samples published in [46, 48, 49, 65, 148].

In Central/South America (shortened to ‘Americas’ in the manuscript), we used 1kG Peruvians (PEL; *N* = 69) and HGDP Mayans (MAYA; *N* = 18) for our present-day populations, and included IBS to represent the European ancestry present in many present-day Americans. We left the ancient samples in one group (ANCA; *N* = 216), and used k=2 for ADMIXTURE (Figure S5). The ancient samples used here were originally published in [149–161]. To construct the PCA we used Human Origins samples published in [46, 48].

#### Power Simulations

To estimate the power of our analysis, we conducted Wright-Fisher simulations of selection for each of the five regions using the model described in Supplementary Note A. For each population, we randomly selected 1000 variants with MAF *>* 0.05, initialized them at their observed frequencies, and simulated their allele-frequency trajectories for 0, 10, 20, 50, or 100 generations with *s ∈ {*0, 0.01, 0.02, 0.05, 0.1*}*. We then conducted the selection scan as for the real data, substituting the simulated frequency for the relevant population and using the same genomic-control and multiple-testing corrections as for the real scan.

#### Comparison with other selection scans

To determine how our results compare to those of other selection scans, we compared our European scan (with the highest power and most other scans to compare) to 4 other scans: [1, 3, 6, 8]. For each set of published scores we intersected the variants tested with those in our scan. Because Kerner et al. only published a list of significant tag SNPs, we also pruned the significant results of the other three, as well as ours, then calculated the enrichment of the hits in each scan in our top N% of peaks. We pruned using Plink’s clumping function with a window of 1Mb and *r*^2^ *>* 0.4 [162]. See Supplementary Note B for further details and discussion.

#### Time Series Simulations

To characterise the performance of bmws-admix we developed a simulation framework allowing us to mimic observed data to varying degrees (Supplementary Note D). We simulated our populations with *k* = 3 ancestry components, since that was the number we used for our best-powered selection scans. We then used a Wright-Fisher framework for each ancestry component independently, allowing for variable *s*, then sampled admixed individuals at each time point to get the overall allele frequencies. We also simulated neutral loci that were strongly differentiated among ancestry components while ancestry proportions changed substantially through time. These simulations produced large changes in pooled allele frequency but no systematic signal of selection with bmws-admix, whereas ancestry-unaware bmws inferred positive selection (Figure D4).

### QUANTIFICATION AND STATISTICAL ANALYSIS

#### A variant-level test for selection

To test for selection while taking into account admixture history, we explicitly model relationships between ancient and modern populations (Figure S15). Specifically, we use admixture proportions to calculate the expected allele frequencies for a variant, and test whether observed frequencies match that expectation [1]. Unlike the previous iteration of this method, we infer admixture proportions in both ancient and modern populations relaxing the assumption that we have samples from unadmixed source populations and more accurately reflecting the fact that ancient populations were themselves products of admixture of previous peoples.

We quantify these relationships in a likelihood framework. The log-likelihood for a given variant under the alternative hypothesis that allele frequencies across populations are unconstrained is

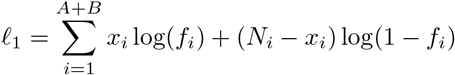

where *A* and *B* are the number of ancient and present-day populations respectively, *x*_*i*_, *N*_*i*_, and *f*_*i*_ = *x*_*i*_*/N*_*i*_ are the observed alternate allele count, total allele counts and allele frequency for the variant in population *i*. To obtain the likelihood under the null hypothesis that allele frequencies are constrained, *l*_0_, we calculate the expected frequencies in our set of populations as:

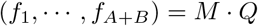

Where

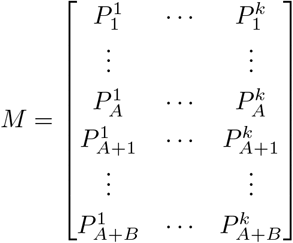

and

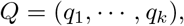

where *M* is a (*A*+*B*) ×*k* matrix containing the admixture proportions for the *A*+*B* ancient and present-day populations for each of the *k* source populations, and *Q* is a vector of length *k* containing the allele frequencies of the variant in each of the source populations. We estimate *M* using ADMIXTURE v1.3.0 [12], after conducting LD pruning (200kb windows, 25-variant step, *r*^2^ *>* 0.4). For the non-European samples, we ran ADMIXTURE in unconstrained mode, across a range of source population numbers from *K* = 2 to *K* = 6 and used the *K* with the minimum cross-validation error. (Within Europe we fixed *K* = 3 because of well-established evidence for three source populations.) Conditional on *M*, we fit *Q* to maximize

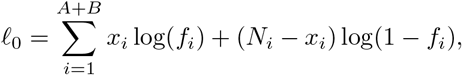

thereby obtaining both the null likelihood and the expected frequencies in our source populations. *Q* is equivalent to the *P*-matrix generated by ADMIXTURE but we fit it numerically for consistency with the calculation of the alternative likelihood. The value of 2(*l*_1_ *− l*_0_) follows a *χ*^2^ distribution with *A* + *B – k* degrees of freedom. We additionally controlled for inflation due to genetic drift or unmodeled ancestry variation using genomic control [163], dividing the *χ*^2^ statistics by a constant so that the median statistic matches the theoretical value, and calculate a P-value based on the 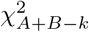 distribution.

We ran this scan for 1,150,640 variants captured by the 1240k reagent commonly used for ancient DNA data [10, 11]. Because we were combining genetic data across data sets, technical artifacts from one dataset could drive false positives in this scan. We therefore removed all variants with FDR *>* 0.1 that did not have at least one (two for Europe) additional variant within 200kb and high LD (*r*^2^ *>* 0.5) also pass that FDR threshold. We calculated LD for this purpose using the 1kG populations corresponding to each region. We then focused on signals passing a genome-wide Bonferroni multiple testing correction with *p <* 5 × 10^*−*8^. We plotted individual significant signals using LocusZoom [164].

For each significant signal, we tested for replication in the other regions both of the lead variants and of the locus more generally. To test if the lead variants replicated we examined their P-values in each other region, using a Bonferroni correction for the number of lead variants. To test for the general locus, we identified the minimum p-value within a megabase of the lead SNP, then used a Bonferroni correction for the number of SNPs tested in the window to determine replication.

It is difficult to calibrate selection tests in the HLA region for a variety of reasons, including high levels of linkage disequilibrium; a different population history than other portions of the genome (stemming from, e.g. the effects of balancing selection), and the fact that high SNP density can lead to spurious visual peaks in genome-wide scans. Therefore, within the HLA, we used a separately calibrated empirical null distribution, and also performed a 50kb window-based Bonferroni correction to control for SNP density (Figure S6).

#### Time series analysis

To analyze time series of allele frequencies while accounting for population continuity and admixture, we extended bmws [2] to incorporate individual-specific admixture loadings. In this section we describe those extensions.

We adopt the convention that *t* = 0 is the oldest modeled generation and *t* = *T* is the present, so all transitions proceed from *t −* 1 to *t*. Let *K* be the number of ancestry components and let *N*_*e*_ = 10,000 be the constant diploid effective population size used in our analyses. For ancestry component *k, F*_*tk*_ *∈ {*0, …, 2*N*_*e*_*}* is the derived allele count and

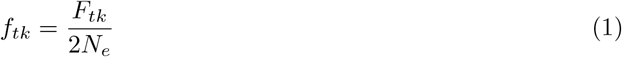

is its frequency. Throughout the manuscript, uppercase *F* denotes an allele count, lowercase *f* denotes an allele frequency, and 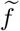 denotes a post-selection frequency.

The coefficient *s*_*tk*_ governs selection during the transition from generation *t −* 1 to generation *t*. Following the original bmws parameterization [2], the derived allele has relative fitness 1 + *s*_*tk*_*/*2. We write its frequency after selection, but before Wright-Fisher sampling, as

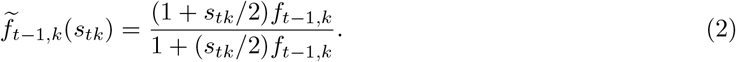

The transition model is

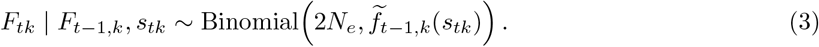

There is no mutation in the model, so loss (*F*_*tk*_ = 0) and fixation (*F*_*tk*_ = 2*N*_*e*_) are absorbing states.

Suppose that we observe *n*_*t*_ alleles in generation *t*. Let *a*_*ti*_ ∈ {0, 1} denote the ancestral or derived state, respectively, of observed allele *i*. A diploid observation can equivalently be represented as two alleles. Let *q*_*tik*_ be its ancestry loading for component *k*, with ∑ _*k*_ *q*_*tik*_ = 1, and let the unobserved ancestry of the allele be *z*_*ti*_ *∈ {*1, …, *K}*. The matrix 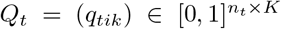 is obtained from ADMIXTURE [12].

The observation model is

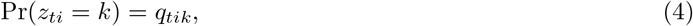

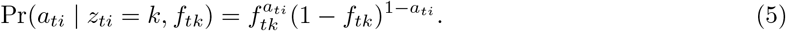

#### Gibbs sampler for selection and allele frequency trajectories

To sample allele-frequency trajectories, latent ancestry assignments, and selection coefficients, we use a Gibbs sampler that alternates among the following conditional updates:

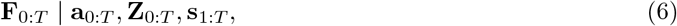

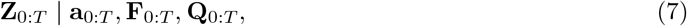

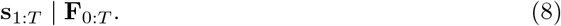

The allele-frequency trajectory is the most difficult of these three quantities to update. In the single-population bmws model, the distribution of *f*_*t*_ was represented by a mixture of beta distributions with point masses at loss and fixation [2]. Directly extending that representation to *K* different ancestry components would require a mixture with (*M* +2)^*K*^ terms. We instead approximate the joint distribution with a conditional sequential Monte Carlo particle filter with ancestor sampling [165, 166]. At generation *t*, particle *j* is a vector of allele counts 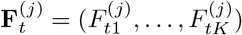 with normalized weight 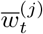. The filtering distribution is approximated by

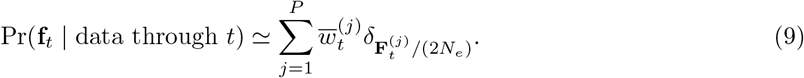

During the Gibbs update, the current **Z**_0:*T*_ and **s**_1:*T*_ are held fixed while the particle filter constructs an updated trajectory **F**_0:*T*_. With a finite number of particles, selecting a trajectory from an ordinary particle filter gives only an approximate draw from the conditional posterior, and therefore cannot be used directly as a proper Gibbs update. Particle Gibbs [167] instead conditions the particle filter on the trajectory retained from the preceding iteration by including that trajectory among the *P* candidates. This construction defines a Markov transition that leaves the desired conditional posterior of **F**_0:*T*_ invariant. We call the retained trajectory the *reference trajectory*; it is an algorithmic bookkeeping device needed to make the sampler correct, and has no biological significance.

At every generation the reference trajectory allele count is fixed to the value retained from the previous generation, whereas the other *P −* 1 particles are free to explore alternative values. At the end of the filter, one of the *P* complete trajectories is selected and becomes the reference trajectory for the next iteration. Before the first iteration, the reference trajectory is initialized from the mean of bootstrap logistic regressions. This initialization improves efficiency but does not alter the target posterior distribution.

For the prior, we place independent Beta(1, 1) priors on the *K* allele frequencies at the oldest generation. Conditional on the current ancestry assignments, let *D*_0*k*_ and *A*_0*k*_ be the numbers of derived and ancestral observations assigned to component *k*. The *P −* 1 non-reference particles are initialized from

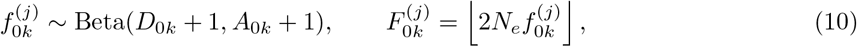

with equal weights, while the remaining particle is set to the oldest state of the reference trajectory.

For each subsequent generation *t* = 1, …, *T*, each of the *P −* 1 non-reference particles selects a parent from generation *t −* 1 with probability proportional to the particle weights. The state of the reference particle at generation *t* is fixed, but its parent is not. We use *ancestor sampling* [166] to select that parent, assigning a previous particle probability proportional to its weight multiplied by the Wright-Fisher probability of transitioning from that particle into the fixed reference state. This allows the retained portion of the trajectory to reconnect to a different, plausible history instead of forcing the entire trajectory from the preceding Gibbs iteration to remain intact. It therefore reduces path degeneracy and improves mixing of the Gibbs sampler. (Here “ancestor” means the parent of a particle in the algorithm, and is unrelated to the biological ancestry components indexed by *k*.)

At each time step, particles are resampled according to their weights to prevent degeneracy. After resampling, we propagate the particles using a proposal that incorporates the observations at generation *t*. Let 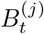 denote the resampled ancestor index of particle *j*, and define *D*_*tk*_ = ∑_*i*_**1**(*z*_*ti*_ = *k*)*a*_*ti*_ and *A*_*tk*_ =∑_*i*_ **1**(*z*_*ti*_ = *k*)(1 *− a*_*ti*_). For a particle with parent 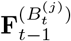, let

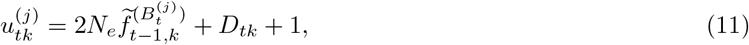

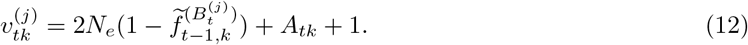

The joint proposal mass is

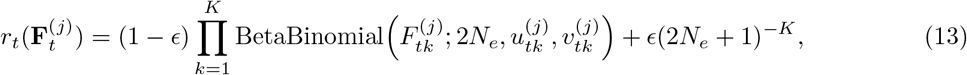

where the second term is a uniform distribution (with *ϵ* = 10^*−*6^) used to ensure absolute continuity of the proposal distribution and improve stability. If generation *t* has no observations, we sample directly from Equation (3) instead.

For a generation with observations, the unnormalized importance weight of particle *j* is

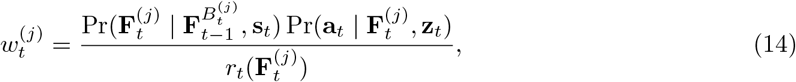

After the final generation, we sample one particle according to 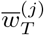 and follow its recorded parents back to *t* = 0. The resulting complete trajectory **F**_0:*T*_ is the updated Gibbs sample, and is used as the reference trajectory in the next iteration.

For the remaining Gibbs updates, conditional on the allele-frequency trajectory, the *z*_*ti*_ are independent and are sampled according to

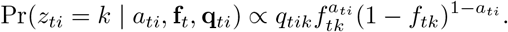

Finally, **s**_*t*_ is conditionally independent of the remaining variables given (**F**_*t−*1_, **F**_*t*_). Its conditional density is the transition probability in Equation (3) multiplied by the regularization prior described below. This density is not a standard distribution, so we update the selection trajectory with the Metropolis-adjusted Langevin algorithm [168]. Each ancestry-specific trajectory is represented by 10 equally spaced linear-spline coefficients.

All estimates used *P* = 5,000 particles, 5,000 Gibbs iterations, *N*_*e*_ = 10,000. We discarded the first 500 iterations and used every fiftieth subsequent state for the posterior summaries.

#### Regularization

In the original `bmws` method, selection coefficient trajectories were regularized via the smoothing penalty 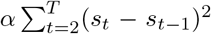 to prevent overfitting, where *α* is a tuning parameter controlling the amount of regularity. As *α → ∞*, the estimate **s** shrinks to a constant value, implying no time-varying selection.

Here we incorporate an additional term that encourages between-population similarity in these trajectories, in recognition of the fact that selective pressures within and among the various ancestral populations likely covaried, at least partially. To model this, we add an additional penalty term of the form

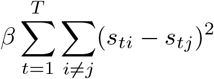

which penalizes the pairwise differences between the selection strength at each time point. As *β → ∞* the model estimates shrink to a single common trajectory (note that this may still be time-varying, depending on *α*), implying that selection does not vary based on ancestry background.

To enable the method to adapt to the level of signal in the data, we adopted a fully Bayesian approach, placing hyperpriors on *α* and *β*. Specifically, we used independent Gamma priors *α ~* Gamma(*a*_*α*_, *b*_*α*_) and *β ~* Gamma(*a*_*β*_, *b*_*β*_), where *a*_*α,β*_ and *b*_*α,β*_ are hyperparameters. Then, *α* and *β* are sampled as part of the Gibbs sampler described above according to standard updates:

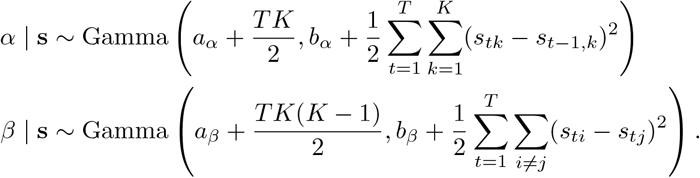

To choose the prior centers, we used an empirical-Bayes approach. We fitted the model over a grid of values for log_10_(*α*_0_), setting *β*_0_ = *α*_0_*/*100 in each fit. Within each fit, *α, β*, and the selection trajectory were sampled during the Gibbs iterations. For each variant, we compared the mean particle-filter log-likelihood across different settings for the prior (Figure S16). Based on this comparison, we used log_10_(*α*_0_) = 1.5, corresponding to *α*_0_ = 10^1.5^, and *β*_0_ = *α*_0_*/*100.

#### Test for polygenic selection

To test for selection across multiple variants contributing to a polygenic trait, we adapted the same pipeline used for the locus-based test. Instead of counting the number of alternate alleles for a single variant across populations, we counted the proportion of trait-increasing alleles across all significant variants identified in a genome-wide association study (GWAS). For population *i, N*_*i*_ is the total number of non-missing (pseudohaploid) genotypes across all individuals and significant variants, and *x*_*i*_ is the total number of genotypes with the trait-increasing allele. The likelihood calculation is then exactly the same as for the single variant test, but we calculate P-values based on a null distribution obtained by permuting the direction of effects and recalculating the likelihood, rather than based on the *χ*^2^ distribution.

We did this for 33 biometric and disease traits analysed in European ancestry individuals: Alzheimer’s Disease [169], Asthma [170], Covid-19 (3 degrees of severity; Freeze 7 of the HGI) [171], Coronary Artery Disease (CAD) [172], Inflammatory Bowel Disease, ulcerative colitis (UC), Crohn’s Disease [173], HDL levels, LDL levels, triglycerides [174], Rheumatoid Arthritis (RA) [175], Type II Diabetes [176], and Waist-hip ratio (all, male, and female, adjusted for BMI) [177], estimated Glomerular Filtration Rate [178]. For the remaining traits we used summary statistics calculated in European-ancestry individuals from the UKBB by the Neale Lab (http://www.nealelab.is/uk-biobank/): Mean corpuscular hemoglobin (MCH), Mean corpuscular volume (MCV), counts of blood cells (platelets, red and white blood cells), Bone Mineral Density, Sitting and Standing Height, Waist Circumference, Hip Circumference, Skin Pigmentation, Tanning, Body Mass Index (BMI), Birth weight and Age at Menopause.

For each trait, we filtered for variants present on the 1240k array, and clumped the association signals using Plink [162]. Clumped SNPs were required to be less than 200 kb apart and have *r*^2^ *>* 0.4. We then counted trait-increasing alleles across all lead 1240k variants with *p <* 10^*−*8^ for each trait. We calculated P-values by permuting the effect direction of each variant included 1,000,000 times, thereby changing which counts were contributing to the trait-increasing alleles. The underlying allele frequencies remained unchanged. We identified traits that showed significant directional and stabilizing selection by calculating 2-sided Bonferroni multiple testing thresholds for *N* traits as 0.025*/N* for directional, and 1 *−* 0.025*/N* for stabilizing selection, where *N* is the number of traits. This choice tends towards being conservative, as we corrected for all traits tested, regardless of their redundancy with others (e.g. we included 3 different metrics of SARS-CoV-2 occurrence).

We also repeated this analysis using 20 traits from GWAS in East Asian-ancestry cohorts: Bone Mineral Density, Body Mass Index, HDL, LDL, Triglyceride levels, Standing Height, Platelet, Red Blood Cell, White Blood Cell counts, and Waist-hip ratio [179]; Asthma, Mean Corpuscular Hemoglobin, Mean corpuscular Volume, Rheumatoid Arthritis, Ulcerative Colitis [180]; Coronary Artery Disease, estimated Glomerular Filtration Rate, Age at Menopause [181]; Type II Diabetes [182]; skin pigmentation [183]. In many cases these GWAS were less powered than the European versions, so for 4 traits with fewer than 40 hits at the stricter threshold (Menopause, UC, Skin Pigmentation, WHR) we lowered the genome-wide significance threshold to 1 × 10^*−*6^ to include enough variants for the selection scan.

## Key Resources Table

| REAGENT or RESOURCE | SOURCE | IDENTIFIER |
| --- | --- | --- |
| **Deposited Data** |  |  |
| Allen Ancient DNA Resource | Reich Lab[35] | doi:10.7910/DVN/FFIDCW |
| 1000 Genomes | 1kG Consortium[37] | N/A |
| Human Genome Diversity Project | HGDP[36] | N/A |
| Alzheimer’s Disease GWAS (EUR) | Kunkle, et al.[169] | doi:10.1038/s41588-019-0358-2 |
| Asthma GWAS (EUR) | Han et al. [170] | doi:10.1038/s41467-020-15649-3 |
| Covid-19 GWAS (EUR) | Niemi et al.[171] | doi:10.1038/s41586-021-03767-x |
| CAD GWAS (EUR) | Aragam et al.[172] | doi:10.1038/s41588-022-01233-6 |
| IBD, UC, Crohn’s Disease GWAS (EUR) | de Lange et al.[173] | doi:10.1038/ng.3760 |
| HDL, LDL, Triglyceride levels GWAS (EUR) | Graham et al.[174] | doi:10.1038/s41586-021-04064-3 |
| Rheumatoid Arthritis (RA) GWAS (EUR) | Ishigaki et al. [175] | doi:10.1038/s41588-022-01213-w |
| Type II Diabetes GWAS (EUR) | Mahajan et al.[176] | doi:10.1038/s41588-022-01058-3 |
| Waist-hip ratio GWAS (EUR) | Pulit et al.[177] | doi:10.1093/hmg/ddy327 |
| eGFR GWAS (EUR) | Stanzick et al.[178] | doi:10.1038/s41467-021-24491-0 |
| UK Biobank | Neale Lab | http://www.nealelab.is/uk-biobank/ |
| East Asian Quantitative Trait GWAS | Chen et al.[179] | doi:10.1016/j.xgen.2023.100436 |
| Biobank Japan GWAS | Sakaue et al.[180] | doi:10.1038/s41588-021-00931-x |
| Biobank Japan GWAS | Kanai et al.[181] | doi:10.1101/2021.09.03.21262975 |
| Type II Diabetes East Asia GWAS | Spracklen et al.[182] | doi:10.1038/s41586-020-2263-3 |
| Skin pigmentation East Asia GWAS | Kim et al.[183] | doi:10.1038/s41467-024-49031-4 |
| **Software and Algorithms** |  |  |
| ADMIXTURE | Alexander et al.[12] | https://dalexander.github.io/admixture/ |
| EIGENSOFT | Patterson et al.[184] | https://github.com/DReichLab/EIG |
| Plinkv1.9b | Chang et al [162] | https://www.cog-genomics.org/plink/ |
| LocusZoom | Boughton et al.[164] | https://my.locuszoom.org/ |
| Julia v1.9.0 | Bezanson et al.[185] | https://julialang.org/ |
| bmws-admix | This Paper | https://github.com/jthlab/bmws_admix |
| Analysis Code | This Paper | https://github.com/colbrall/eas_selection |

## Data and Code Availability

Selection scores and P-values are available through Zenodo (https://doi.org/10.5281/zenodo.18166002), and all genotype data are previously published and publicly available. Scripts for parsing data files and running polygenic and AF-based analyses are available from https://github.com/colbrall/eas_selection. bmws-admix analysis and simulation code are available from https://github.com/jthlab/bmws_admix.

## Acknowledgments

We thank Melinda Yang, Pontus Skoglund, Mary Lauren Benton, Sarah Fong, Ziyue Gao, and members of the Gao and Mathieson labs for helpful discussion and feedback.

## Author Contributions

L.L.C., J.T. and I.M. conceived and designed the study and wrote the manuscript. J.T. implemented the time series analysis, L.L.C. conducted all other analyses. All authors edited and approved the manuscript.

