## Supplementary Material for "Global patterns of natural selection inferred using ancient DNA"

### Supplementary Text

#### A Power analyses

We conducted power simulations in order to better understand what could be driving the differences between the number of significant hits in different regions for the AF-based test. While variable sample sizes would likely influence power, the varying demographic histories and our ability to model them using ADMIXTURE would also be expected to have an influence. We therefore simulated selection using the Wright-Fisher model for random variants in each population in all five regions. We required the MAF of each variant is greater than 0.05, and for each variant set the initial frequency at observed frequency. We did this across a range of selection strengths (0-0.1), and for 1000 random variants in each population, then combined for each region to see how many came up as significant.

As expected, power to detect selection increased with the strength of selection, and was higher in regions with higher sample size (Figure A1). Interestingly, power does indeed seem to decrease with more complicated demography, with the maximum in Africa being about half that in the Americas, despite similar sample sizes. It is also notable that, despite the much larger sample size, even Europe asymptotes at about 70% power. This seems to be primarily driven by how much of the allele frequency trajectory we observe; power is lower for variants that are already at high frequencies at the start of the time series.

Together, this suggests that the difference in apparent selection between regions is likely explained by power differences, influenced potentially both by overall sample size and by the extent to which demographic history is captured by the ADMIXTURE analysis of the samples. To illustrate this, we ran two analyses downsampling the ancient European samples– A) randomly downsampling the whole region to match the overall sample size we had in East Asia, and B) downsampling the individual populations so that they matched the population sizes in East Asia. For A) this resulted in 236 individuals, with 6 HG, 41 EF, 45 EF-ADMIX, 98 BA-N, and 46 BA-S samples, while B had 425 individuals with 23 HG, 94 EF, 103 EF-ADMIX, 102 BA-N and 103 BA-S. In both cases we retained the 4 present-day populations, each with approximately 100 samples.

In both analyses, 2 loci passed a multiple testing correction, despite the overall sample size being identical to that of East Asia where we found four significant loci (Figure A2A-B). This suggests that sample size is the most important factor determining power, although results can be sensitive to the exact composition of the sample. In particular, although we also have hundreds of ancient samples in South Asia and Africa, we do not find any genome-wide significant loci. We hypothesize that this is because the ancient samples do not capture as much of the demographic history of the region as the European and East Asian samples.

We believe this is why the scans in East Asia and the Americas were more successful than in South Asia and Africa, despite similar sample sizes. In both former cases the ADMIXTURE and samples were able to capture known demographic history, reflecting specific known admixture dynamics (northern and southern mainland, and Japan in East Asia, more recent European admixture in the Americas). On the other hand South Asia only had ancient samples from the northern part of the region, and Africa has a deep and complex genomic history that is not well represented with a limited sample size compared to populations that were subject to the out-of-Africa bottleneck. As we increase the sample sizes and our knowledge of demographic history in each region, we will be better able to represent ancient groups (e.g. separating out Central and South American populations).

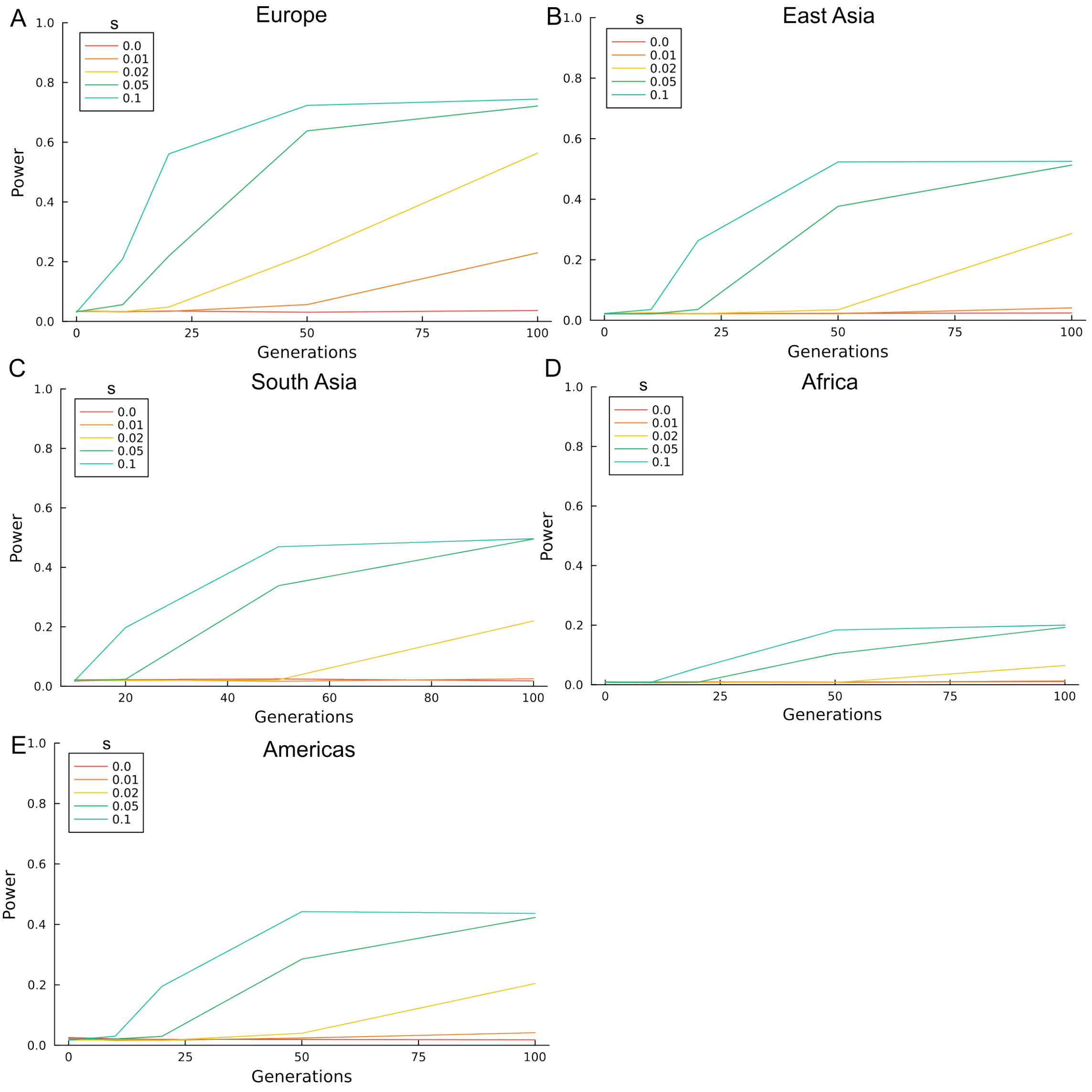


Supplementary Figure A1: Power curves based on Wright-Fisher simulations.
We simulated selection for 1000 random variants in each population in A) Europe, B) East Asia, C) South Asia, D) Africa, E) the Americas. We did this for s = 0, 0.01, 0.02, 0.05, 0.1, and from 0 to 100 generations.

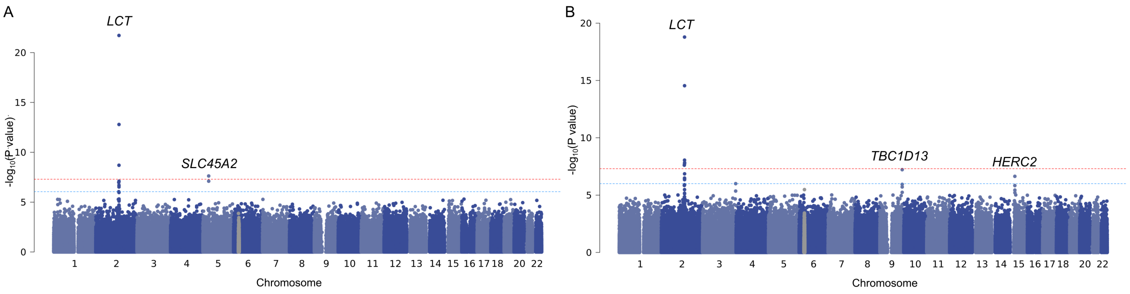


Supplementary Figure A2: European locus-based scans with downsampled ancient groups.
A) Downsampling randomly to 236 ancient individuals overall. B) Downsampling such that the smallest ancient group had 23 samples, while the other four each had approximately 100.

### B Comparison to other European selection scans

While this is the first aDNA-based scan in most of the regions we study here, Europe has much larger availability of aDNA and correspondingly has been studied for selection in similar ways before. We therefore compared our results to both the scan run with the original method modified here [1] and with a recent alternative allele frequency approach [2]. We also compared our results with a study based on local ancestry [3], and with an ABC simulation-based method [4].

In intersecting our results with these other scans, we found broad agreement, with the top hits of each other scan being enriched in our top hits (Figure B1A). Furthermore, this enrichment increased as we were stricter in which hits were considered the "top" ones. It was also notable that the consistently most-enriched results were those from Mathieson 2015, the direct predecessor to this study, while the Kerner et al. with the most distinct methodology were the least enriched (although we lacked access to their full summary statistics, so had to limit ourselves to just the variants they identified as significant).

Despite the broad agreement in patterns of selection across loci, we found the drastic difference in number of significant peaks identified in these studies (ranging from a dozen to hundreds) striking. We particularly focused on Akbari et al., which reported 347 independent significant loci. While some of that increase is due to sample size (about 3000 additional ancient samples), the main factor is their less-stringent genome-wide significance threshold, determined by identifying enrichment of selection signals in GWAS hits, which effectively does half the amount of correction in genomic-control than we implement here. Indeed, if we amend our lambda correction to a similar level we identify 1050 independent loci (Figure B1B).

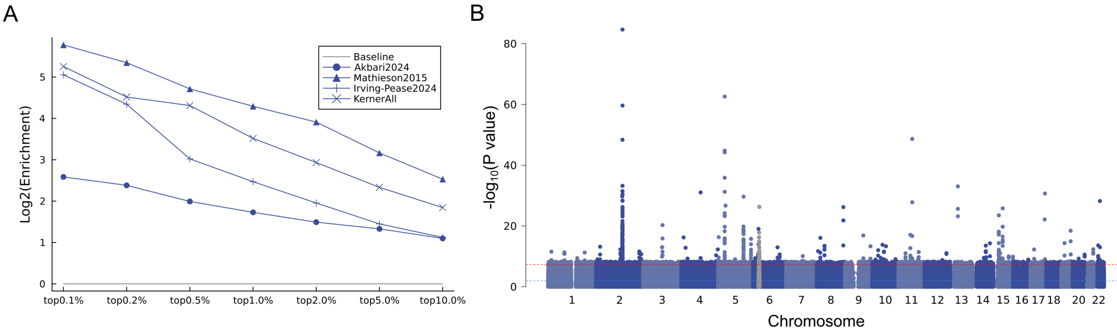


Supplementary Figure B1: Comparison with other European aDNA selection scans.
A) Enrichment of top results from other scans in our top N% hits in Europe. B) Results of our European scan using a GC correction analogous to that used by Akbari et al (lambda = original_lambda*(2.78/5.26)).

### C Discussion of other significant loci

***ABI3BP***; Europe. Lead SNP is in an intron of *ABI3BP*, and is primarily an eQTL for *PDCL3P4*, a pseudogene. *ABI3BP* itself is not well characterized.

***CABP7***; Europe. The lead SNP is in an intron of *ZMAT5*, and is a convincing eQTL for *NIPSNAP1*, a mitochondrial protein with possible involvement in inflammation [5, 6]. It is also an eQTL for *MTMR3*, which is near a GWAS hit for IBS [7].

***CARD8***; Europe, replicated in the Americas and East Asia. The lead SNP replicates in East Asia, and this haplotype is an eQTL for many genes in the locus, in many different tissues, particularly *ODAD1*, and an sQTL for *CARD8* [8]. *ODAD1* is tied to primary ciliary dyskinesia (recurrent lung infections), and is primarily expressed in the lung [9, 10]. *CARD8* is a inflammasome-forming gene with human-specific HIV-sensing functionality [11], as well as wider involvement in inflammatory pathways via different isoforms [12]. The signal in the Americas is much less clear, but does not involve this haplotype.

***CFAP58***; the Americas. The lead SNP is an eQTL for *GSTO1*. *CFAP58* is involved in sperm morphology and development [13], while *GSTO1* is involved in inflammatory pathways and has been associated with severity of symptoms in response to high arsenic exposure [14–17].

***COBLL1***; East Asia, replicated in South Asia. This haplotype is an eQTL for *SCN2A* in cerebellum tissues, and *COBLL1* in testis, as well as *CSRNP1* in skin [8]. *SCN2A* is associated with GWAS traits related to autism and seizures, as well as alcohol use disorder, while *COBLL1* has GWAS hits related to WHR, BMI and lipoproteins, and has variants that modulate obesity [7, 18, 19]. The South Asian peak is an eQTL for *SLC38A11* and *COBLL1*.

***EGFLAM***; Europe, replicated in the Americas. The lead SNP is in an intron of *EGFLAM*, but isn't an eQTL. There are some nearby immune GWAS hits [7], but no obvious functional connection. Not a very clear peak in either region, so difficult to interpret.

***HEATR6***; East Asia. While the lead variant at *HEATR6*, rs2333656, is an eQTL for *HEATR6* and several nearby RNA genes, we do not believe *HEATR6* is a likely target of selection, given its broad expression and lack of known function. However, another variant on the same haplotype (rs345166, $p = 4.1 \times 10^{-}7$), is an eQTL for *CA4*, a zinc-metalloenzyme highly expressed in the lung, colon, adipocytes, and whole blood [8, 20]. Along with other members of the CA gene family, *CA4* catalyzes the hydration of carbon dioxide and may play a role enhancing wider metabolism via interactions with *MCT* transporters [21, 22].

***HS3ST1***; Europe. This peak is 650kb from the gene, not near anything else, and isn't an eQTL. *HS3ST1* is important for heparan sulfate biosynthesis [23]. There are nearby GWAS associations with Alzheimer's Disease, ALS, HDL cholesterol, and BMI [7].

***IRX2***; Europe. This SNP is a long way from both *IRX1* and *IRX2*, with nothing particular around it. There are some GWAS associations with Waist and hip circumference and height [7], but no obvious functional explanation.

***KCNIP1***; Europe. The lead SNP is in an intron of *KCNIP1*, but is not an eQTL for anything. There is a nearby GWAS hit for HIV viral load and some ALL associations, but no obvious link to this variant or gene [7]. *KCNIP1* may be involved in Type II Diabetes [24, 25].

***KHDRBS3***; Europe, replicated in the Americas. The lead SNP is not an eQTL, and is nearly a megabase from this gene, which doesn't have a big depth of literature about it. There is a Bone Mineral Density GWAS haplotype in the general vicinity, but not much else [7]. Signal in the Americas is independent, but similarly opaque.

***MIF***; Americas. Macrophage migration inhibitory factor [26]. The lead SNP is an eQTL for the antisense transcript of *MIF* [8], and we hypothesize that it may regulate *MIF* itself, at least in some conditions (GTEx does not test expression under immune challenge, for example.).

***MXRA7***; Europe. The derived allele of the lead SNP at *MXRA7* was less frequent than expected, primarily in the British population. This haplotype is an eQTL for several genes in the region, though *MXRA7* is significant in the most tissues. The derived allele actually decreases expression of the gene in most tissues, though not all. The gene is involved in macrophage differentiation [27], so potentially associated with host defense.

***NAT2***; Europe. Previously identified locus [28–30], although a different haplotype than previously identified, and the mechanism of selection is unknown. NAT2 interacts with many compounds found in both diet and the environment [31]. The peak is located in an intron of *PSD3*, but is not a known eQTL for anything.

***NUP85***; Europe. The peak at *NUP85*, which is involved in viral response and immune cell migration

[32–34] was previously identified in a time-series-based scan of selection on gene expression in ancient Britons [35], though our results here suggest this is a more general European signal.

***ROR2***; Europe, replicated in Africa. The lead SNP in Europe is an eQTL for *ROR2* and *SPTLC1*. *ROR2* is involved in cartilage and growth plate development, as well as Brachydachtyly type B [36]. There is a nearby GWAS association for height, and BMI [7]. *SPTLC1* is a key enzyme in sphingolipid biosynthesis [37]. The lead SNP in Africa is very rare in Europe, and is almost a megabase away; however this locus has been previously seen in a PBS-based selection scan in Khoe-San [38].

***SLC22A4***; Europe. Previously identified signal [39, 1, 3] near the ergothioneine transporter gene, but also associated with IBD and asthma so potentially related to either diet or immunity.

***UFM1***; Europe. The only eQTL on this haplotype is for *TRPC4*, a cation channel with roles in many processes and is particularly highly expressed in the uterus [8]. *UFM1* is a modifier similar to ubiquitin, and is tied to cancer progression [40].

***ZSCAN31***; Europe (East Asia). The lead SNP is in the intron of this gene, and is an eQTL and sQTL for several nearby genes in many tissues. The nearest GWAS hits are for MDD, Covid-19 and staph aureus infections, pork/oily fish diet, and gas reflux [7]. Most of the genes in the region are understudied zinc fingers, so a functional tie to any of these is not possible. This peak is just outside the MHC region.

### D bmws-admix simulations

To characterize the behavior of bmws-admix, we simulated three ancestry components over approximately 10,000 years (334 generations, assuming 30 years per generation). At the oldest generation, the derived allele had frequency 0.10 in each component. We then simulated an independent Wright-Fisher trajectory for each component with diploid effective population size $N_e = 10{,}000$. For each sampled individual, we first drew the ancestry of the observed allele according to that individual's ancestry proportions and then drew its derived or ancestral state using the contemporaneous allele frequency in that component. All observations were pseudohaploid.

We considered three sampling and admixture scenarios:

1. We sampled 30 individuals every five generations and an additional 100 present-day individuals. Individual ancestry proportions were drawn from a symmetric Dirichlet distribution with concentration parameters $(0.01, 0.01, 0.01)$, producing individuals with almost entirely one ancestry component.

2. We used the same sampling times and sample sizes, but drew ancestry proportions from a Dirichlet$(1, 1, 1)$ distribution, producing entirely admixed individuals with ancestry proportions distributed uniformly over the three-component simplex.

3. We matched the sampling times, sample sizes, and individual ancestry proportions in the European data set.

For each scenario, we varied the strength, timing, and ancestry specificity of selection. Selection either acted throughout the trajectory or during a specified interval, and either had the same coefficient in all three components or acted in only one component; the remaining components had $s = 0$. We simulated 10 replicate data sets for every combination shown in Figures D1–D3. Each replicate was analyzed with 5,000 particles and 2,002 Gibbs iterations (fewer than real analysis for computational time); we discarded the first 500 iterations and retained every twentieth subsequent state. In scenario 1, bmws-admix can detect patterns affecting every component equally down to $s = 0.02$ when selection is transient, or $s = 0.01$ when it is consistent (Figure D1A-D). When selection only impacts one component this can still be detected down to quite small values of $s$ (Figure D1E-H), although in cases where the selected allele fixes, the method struggles to capture the correct magnitude of selection for the whole time span, with the inferred $s$ dropping towards zero.

In scenario 2, where every individual is admixed, bmws-admix is again capable of detecting selection affecting all three components equally (Figure D2A-D),and can detect selection when it impacts only one component when it's strong (Figure D2E-H). However, it doesn't necessarily capture the full trajectory, and it is not obvious how selection could affect only one component of ancestry in a real population where all individuals are admixed.

Finally, we therefore also simulated scenario 3, with the sampling and admixture seen in the European ancient data. Once again it could identify selection affecting all 3 components (Figure D3A,B), but struggled when weak selection affects only 1 (Figure D3C,D). As a general rule in the latter case, scenarios 2 and 3 had a narrow range of s that they could mostly detect, but tended to miss small values of s ($< 0.05$) entirely, and underestimate s when it approached 0.1.

To test directly whether changing ancestry proportions can produce spurious evidence of selection, we simulated 10 replicate neutral data sets ($s = 0$) over 100 generations. The three ancestry components had initial derived allele frequencies of 0.05, 0.50, and 0.95. Mean ancestry proportions changed linearly from $(0.80, 0.10, 0.10)$ 100 generations ago to $(0.10, 0.10, 0.80)$ at present, and individual ancestry proportions were sampled around these means. We sampled 100 pseudohaploid observations every five generations. This change in ancestry produced a median increase of 0.617 in the pooled allele frequency despite the absence of selection (Figure D4A,B). We analyzed each replicate both with bmws-admix, using the individual ancestry proportions, and with an ancestry-unaware, single-component bmws model.

The bmws-admix estimates showed no systematic signal of selection: the median selection coefficients across time and replicates were $-0.010$, $-0.003$, and 0.007 for the three components, and 26 of the 30 corresponding 95% posterior intervals included zero (Figure D4C). In contrast, the ancestry-unaware model interpreted the pooled frequency change as positive selection in all 10 replicates, with a median time-averaged selection coefficient of 0.049 (Figure D4D). These simulations show that explicitly modeling ancestry separates allele-frequency changes caused by admixture from changes within ancestry components caused by selection.

Overall we find that bmws-admix works well when selection coefficients are the same across ancestries; in which case the different ancestral estimates for $s$ should be regarded as distinct estimates of the same underlying parameters. It can also detect differing selection coefficients across ancestry components, though has less power. Overall it is relatively robust to differing ancestry patterns. In particular, if selection and ancestry both vary with geography then the average selection coefficient experienced by each ancestry might differ; in this case bmws-admix will effectively estimate the average selection coefficient experienced by individuals in the sample. It is also worth noting that once the derived allele fixes, selection can no longer be reliably detected; distinguishing between that scenario vs a transient selective pressure requires examining allele frequencies as well as the selection trajectories.

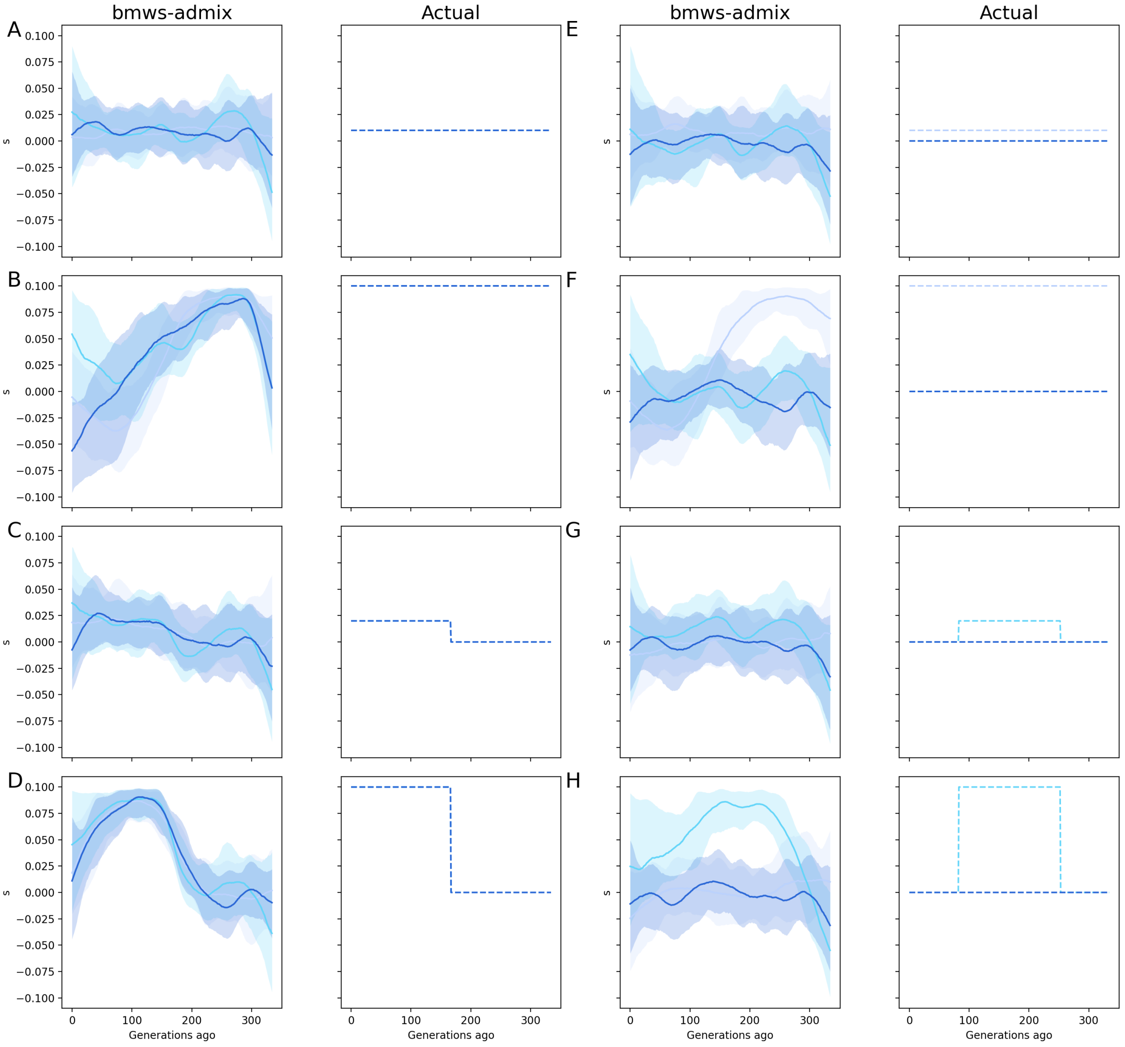


Supplementary Figure D1: Comparison of simulated selection and results from bmws-admix for simulations including unadmixed individuals

We simulated equal selection for all three components for A) $s = 0.01$ and B) $s = 0.1$ across the full time span, and C) $s = 0.02$ and D) $s = 0.1$ for the most recent half of the time. We also simulated with only one admixture component under selection for E-F) the full timespan and G-H) the middle half.

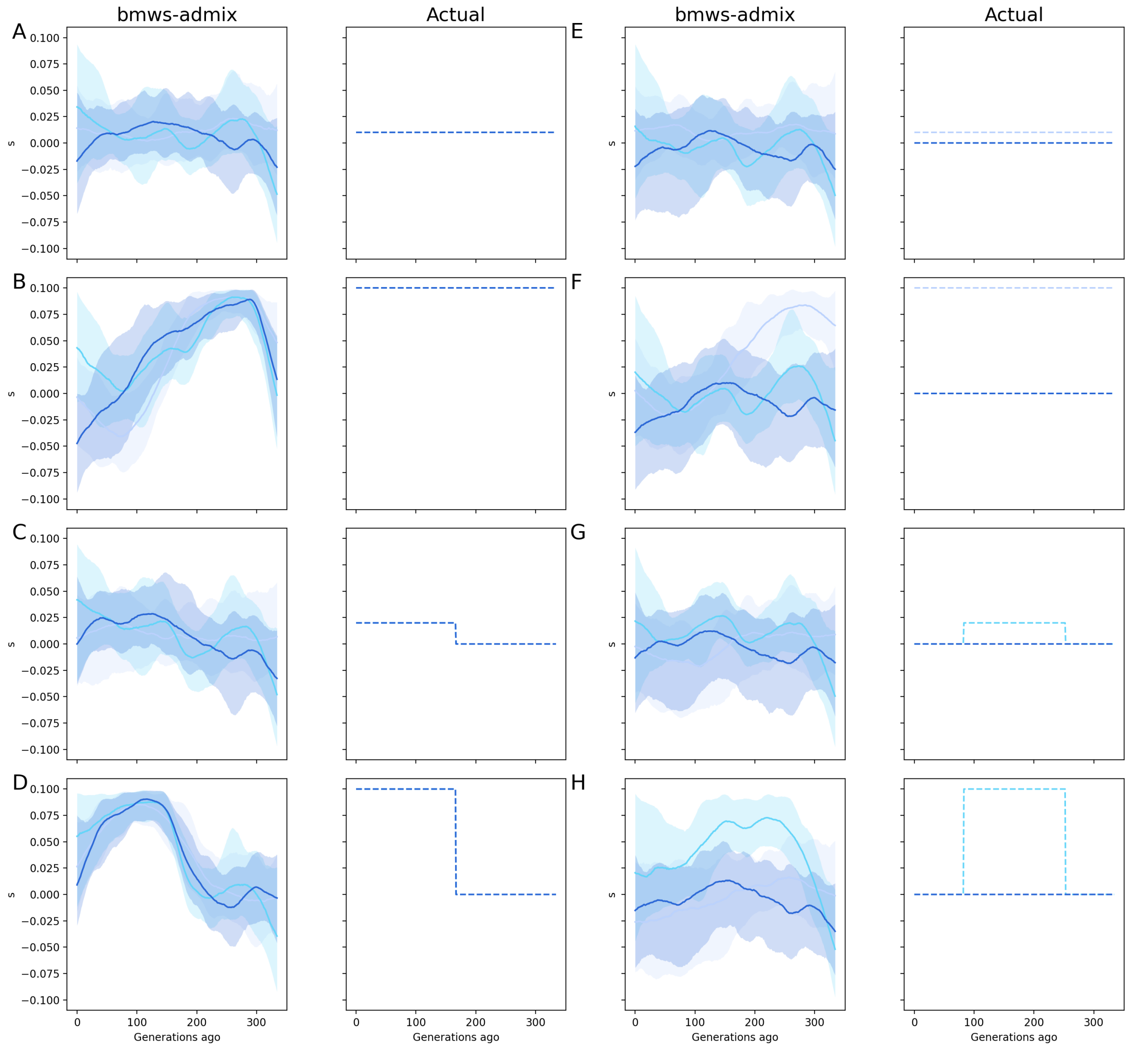


Supplementary Figure D2: Comparison of simulated selection and results from bmws-admix for simulations for entirely admixed individuals

We simulated equal selection for all three components for A) $s = 0.01$ and B) $s = 0.1$ across the full time span, and C) $s = 0.02$ and D) $s = 0.1$ for the most recent half of the time. We also simulated with only one admixture component under selection for E-F) the full timespan and G-H) the middle half.

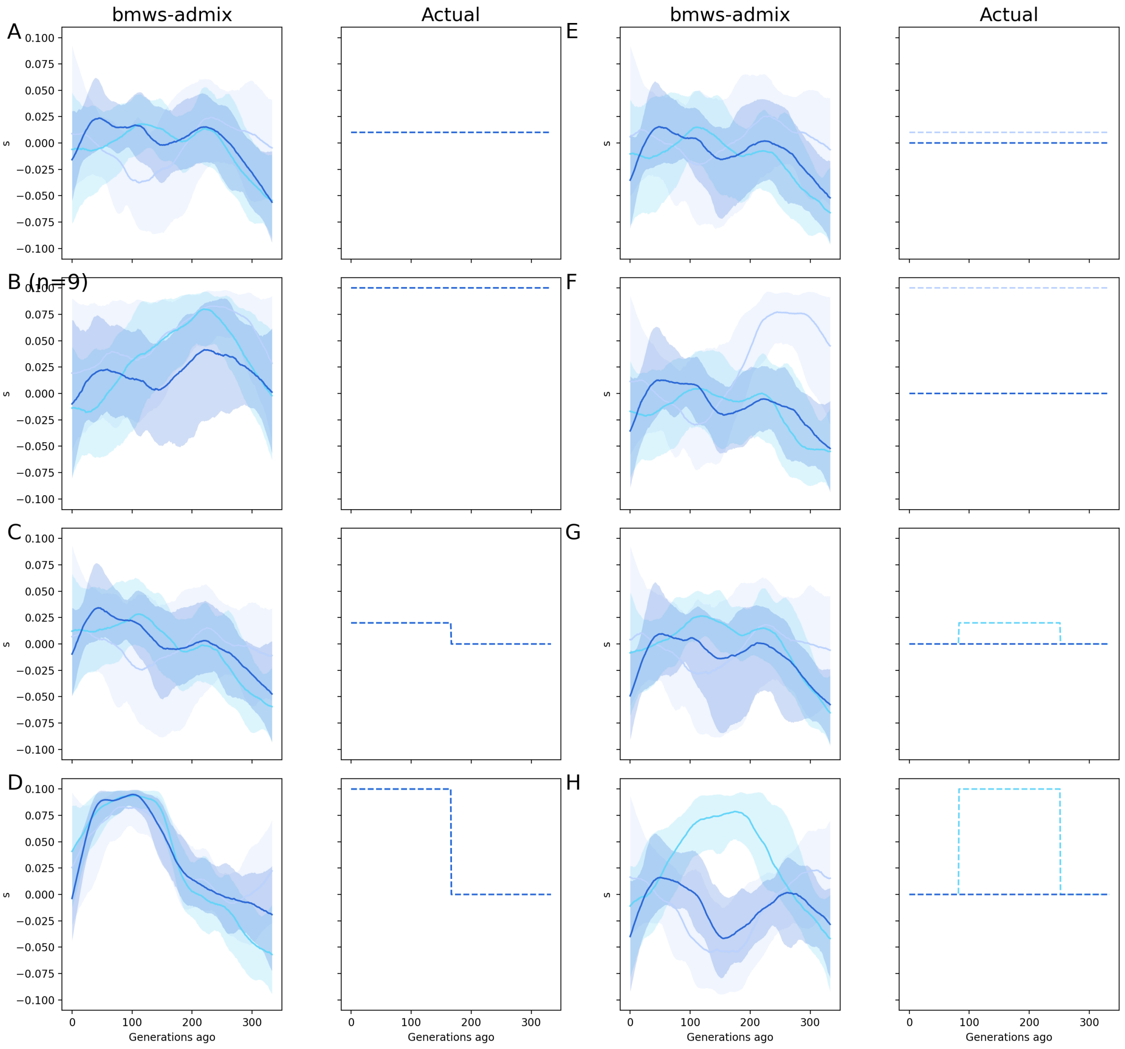


Supplementary Figure D3: Comparison of simulated selection and results from bmws-admix for simulations matching the timing, sampling, and admixture proportions of Europe
We simulated equal selection for all three components for A) $s = 0.01$ and B) $s = 0.1$ across the full time span, and C) $s = 0.02$ and D) $s = 0.1$ for the most recent half of the time. We also simulated with only one admixture component under selection for E-F) the full timespan and G-H) the middle half.

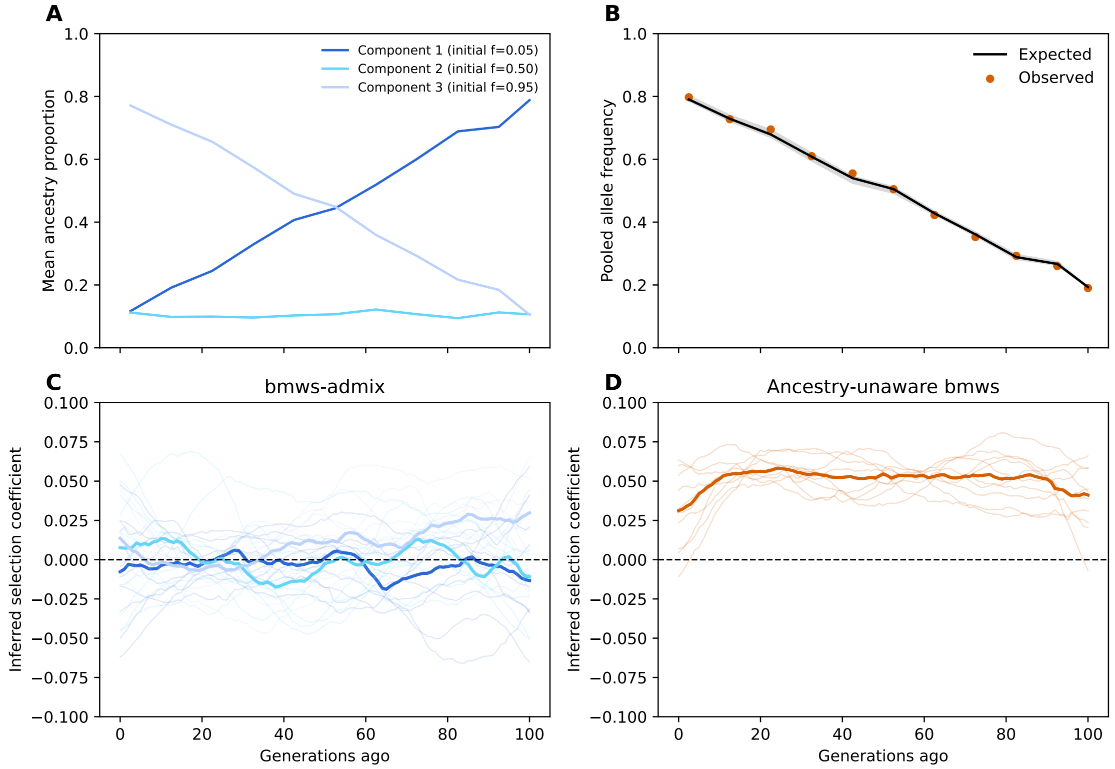


Supplementary Figure D4: Allele frequency change at a neutral locus caused by a shift in ancestry proportions. A) Mean ancestry proportions over time. The three ancestry components had initial derived allele frequencies of 0.05, 0.50, and 0.95. B) Expected and observed pooled allele frequencies. The black line is the median expected frequency, the gray band spans the 5th–95th percentiles across 10 replicates, and orange points are the median observed frequencies. C) Selection trajectories inferred by bmws-admix for each ancestry component. Thin lines show the posterior median for each replicate and thick lines show the median across replicates. D) Corresponding trajectories inferred by ancestry-unaware bmws. All simulations were neutral.

### Supplementary Figures

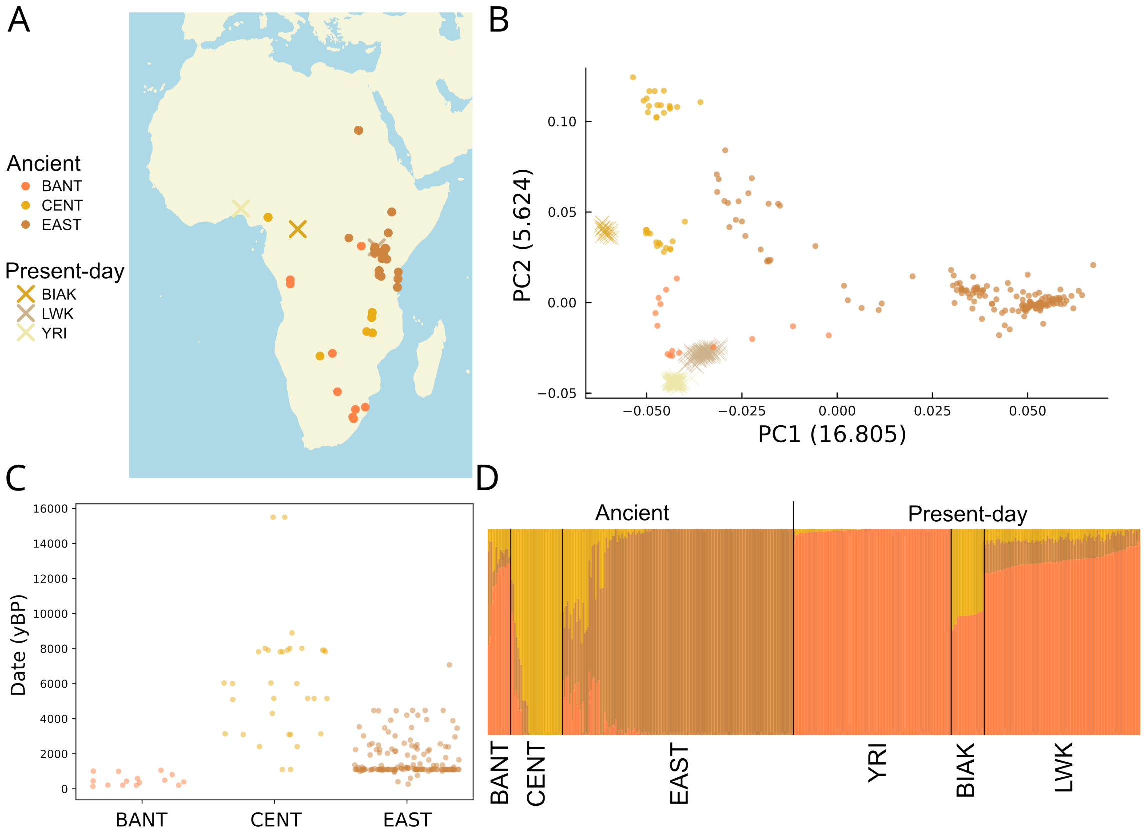


Supplementary Figure S1: Summary of the African groups used in this study.
A) Map of samples (for ancients) and collection points (for present-day). B) PCA with the 3 ancient and 3 present-day groups constructed using African samples from Human Origins dataset. C) Age distributions of ancient groups. D) ADMIXTURE plot of the ancient groups and present-day populations, run with $K = 3$. We took the mean (weighted by coverage) of these admixture proportions for each population to use for the selections scans. BANT= Bantu; CENT = Central African; EAST = East African; YRI = Yoruba, LWK = Luhya, BIAK = Biaka.

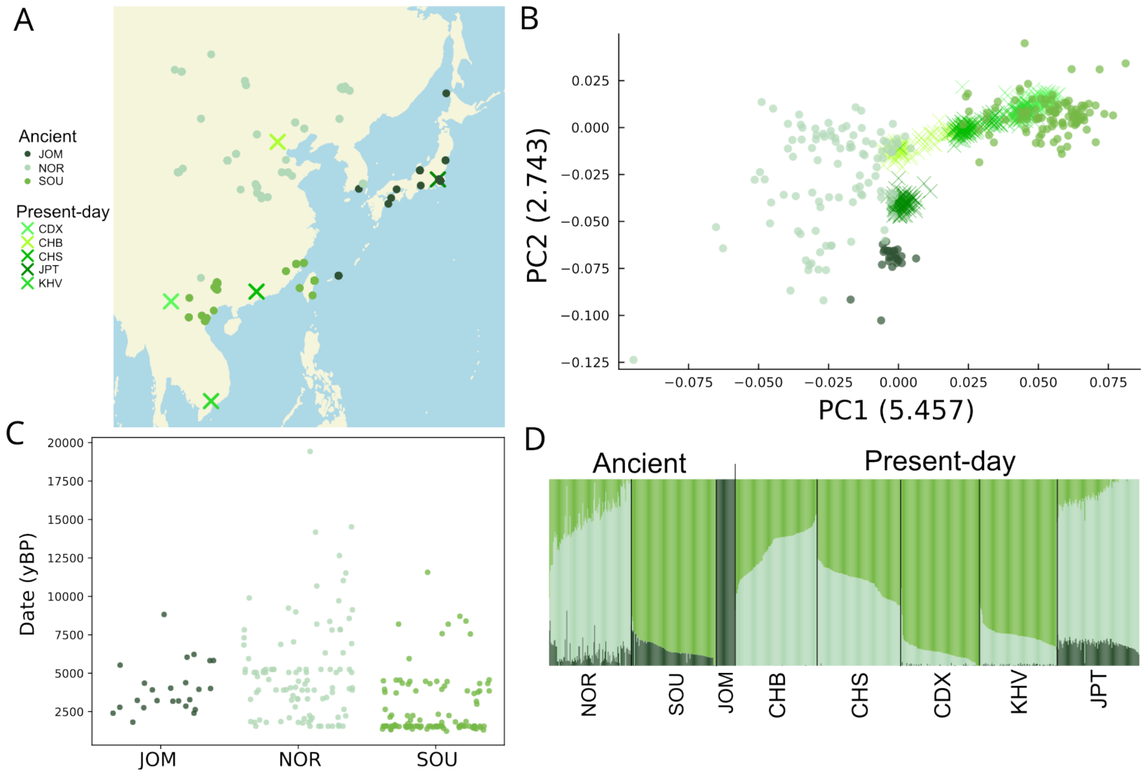


Supplementary Figure S2: Summary of the East Asian groups used in this study.
A) Map of samples (for ancients) and collection points (for present-day). B) PCA with the 3 ancient and 5 present-day groups constructed using Asian samples from Human Origins dataset. C) Age distributions of ancient groups. D) ADMIXTURE plot of the ancient groups and present-day populations, run with $K = 3$. We took the mean (weighted by coverage) of these admixture proportions for each population to use for the selections scans. NOR = Northern Mainland; SOU = Southern Mainland; JOM = Jomon; CHB = Han Chinese (Beijing); CHS = Han Chinese (South); CDX = Dai; KHV = Vietnamese; JPT = Japanese.

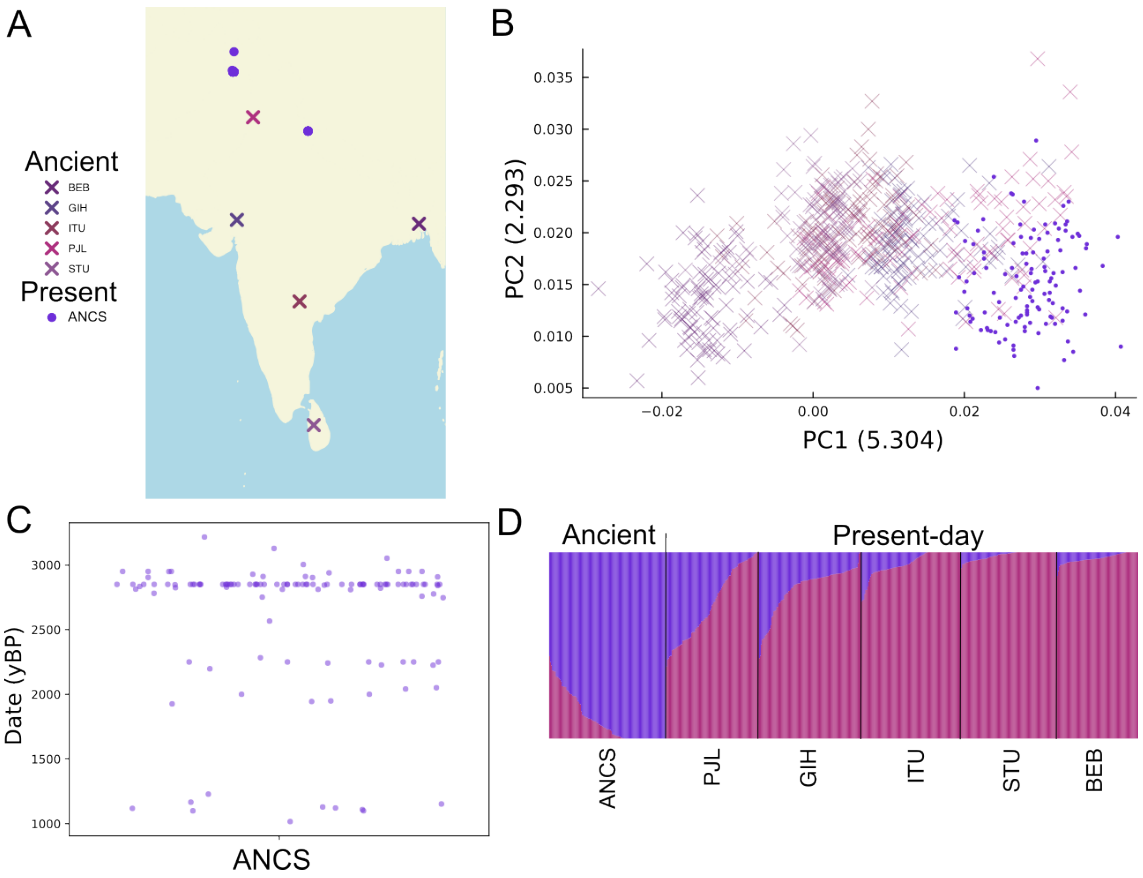


Supplementary Figure S3: Summary of the South Asian groups used in this study.
A) Map of samples (for ancients) and collection points (for present-day). B) PCA with the 1 ancient and 5 present-day groups constructed South Asian samples from the Human Origins dataset. C) Age distributions of ancient group. D) ADMIXTURE plot of the ancient groups and present-day populations, run with $K = 2$. We took the mean (weighted by coverage) of these admixture proportions for each population to use for the selections scans. ANCS = Ancient; PJL = Punjabi; GIH = Gujarati; ITU = Telugu; STU = Tamil; BEB = Bengali.

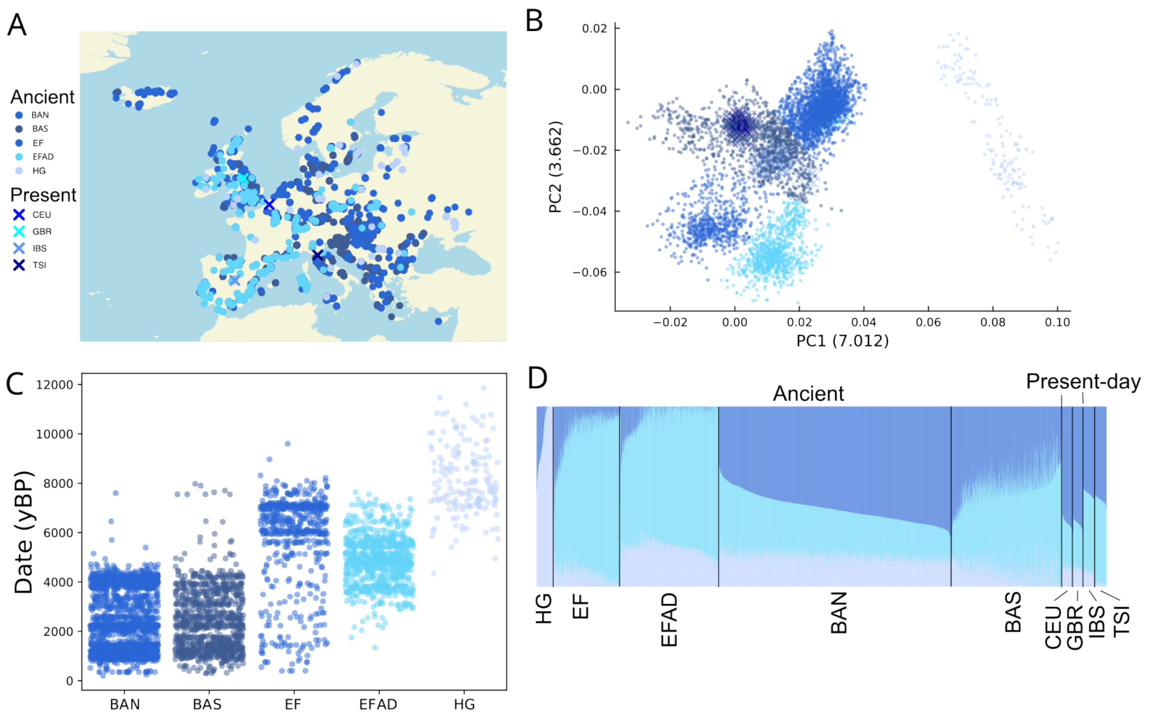


Supplementary Figure S4: Summary of the European groups used in this study.
A) Map of samples (for ancients) and collection points (for present-day). B) PCA with the 5 ancient and 4 present-day groups constructed using Non-Finnish European samples from Human Origins dataset. C) Age distributions of ancient groups. D) ADMIXTURE plot of the ancient groups and present-day populations, run with $K = 3$. We took the mean (weighted by coverage) of these admixture proportions for each population to use for the selection scans. HG = Hunter-gatherer; EF = Early Farmer; EFAD = Early Farmer (Admixed); BAN = Bronze Age (North); BAS = Bronze Age (South); CEU = CEPH Northwestern European; GBR = British; IBS = Spanish; TSI = Tuscan.

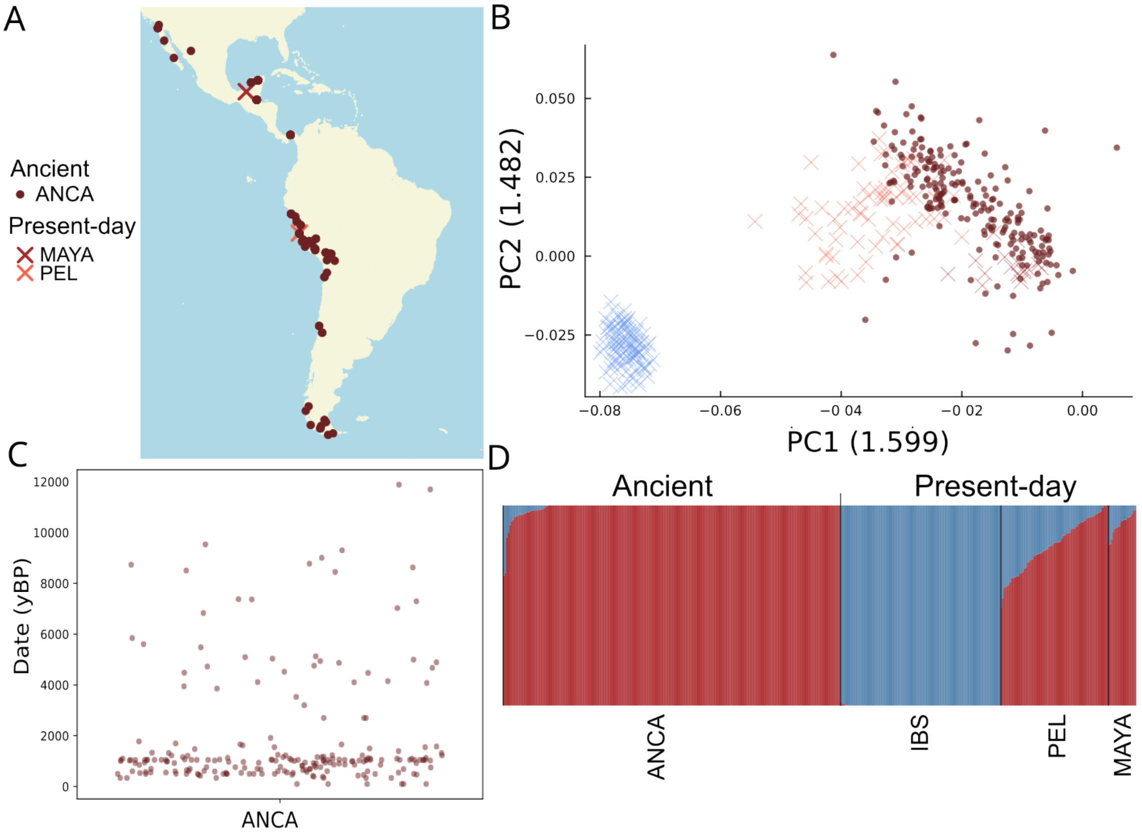


Supplementary Figure S5: Summary of the American groups used in this study.
A) Map of samples (for ancients) and collection points (for present-day; excluding IBS, which is marked on the Europe map in Supp. Fig. S4. B) PCA with the 1 ancient and 3 present-day groups constructed using Asian samples from Human Origins dataset. C) Age distributions of ancient groups. D) ADMIXTURE plot of the ancient groups and present-day populations, run with $K = 2$. We took the mean (weighted by coverage) of these admixture proportions for each population to use for the selections scans. ANCA = Ancient; IBS = Spanish; PEL = Peruvian; MAYA = Mayan.

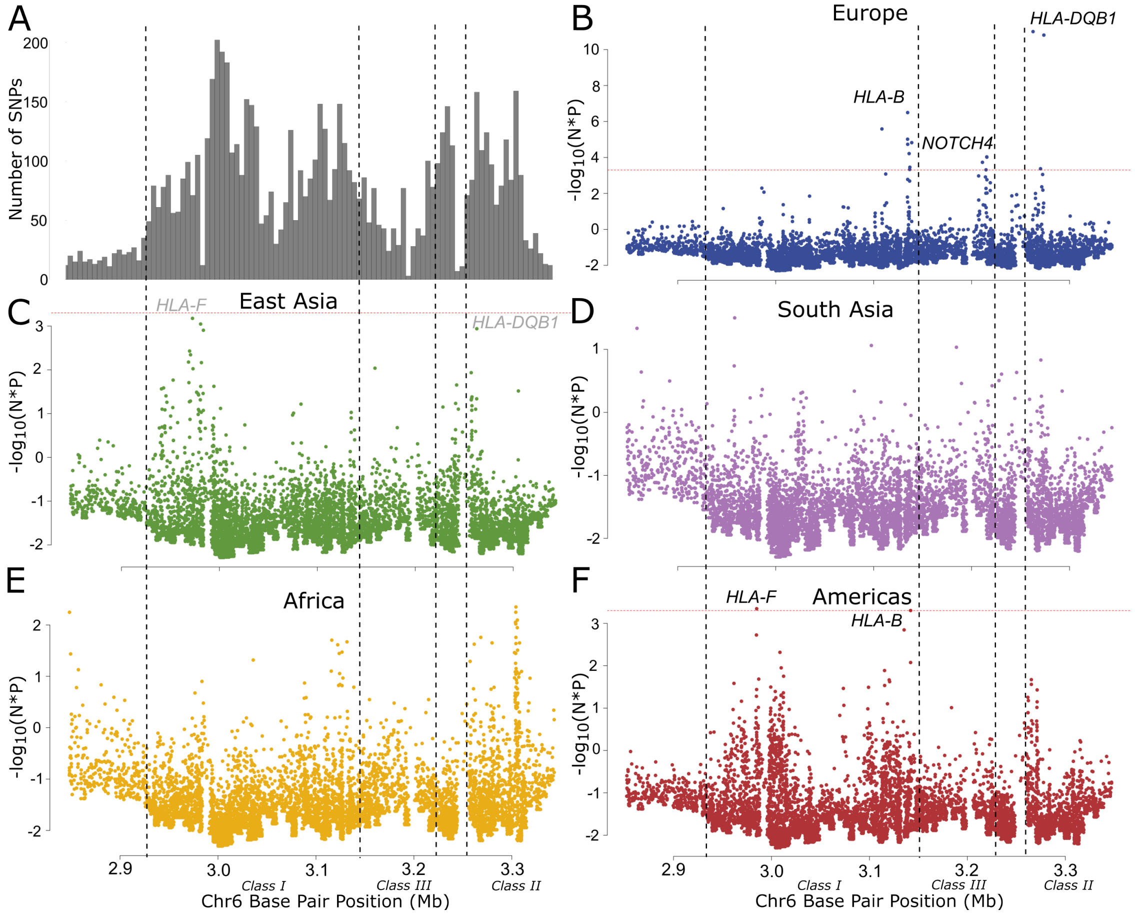


Supplementary Figure S6: Manhattan plots corrected for SNP density focused on the MHC. We corrected for A) the distribution of SNPs across the MHC region in B) Europe, C) East Asia, D) South Asia, E) Africa, and F) the Americas. Each P-value in the manhattan plots is corrected by multiplying P by the number of SNPs in a tiled 50kb window, then the multiple testing threshold (red dashed line) corrects for the number of bins across the HLA. Dashed black boxes indicate the approximate locations of Class I, II and III HLA genes. Significant peaks are labelled in black with their nearest gene; grey labels indicate nominally-replicating sites.

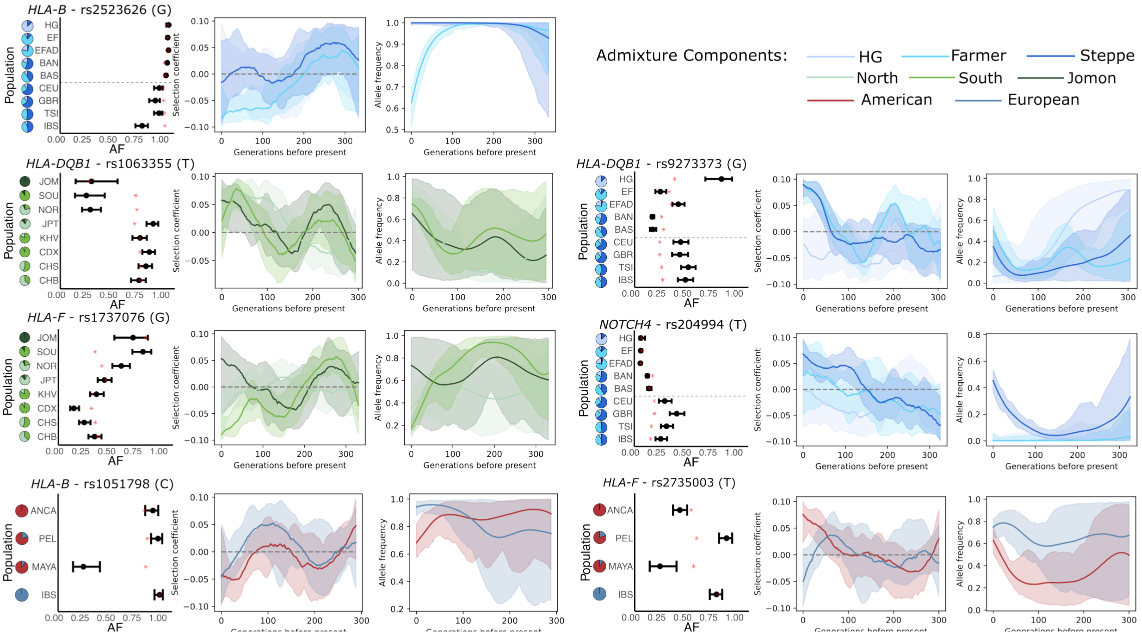


Supplementary Figure S7: Locus plots for significant and replicating peaks in the HLA region. From left to right, we plotted Observed AFs, selection coefficient, and inferred AF trajectories. We plotted the information for the derived allele of the lead SNP in each peak. In the frequency plots, the error bars represent the 95% confidence intervals calculated using the Agresti-Coull method, while the red asterisk indicates the fitted frequency from the admixture model and the dotted line separates ancient (top) from present-day (bottom) groups. The pie charts indicate the average proportion of each admixture component for each population.

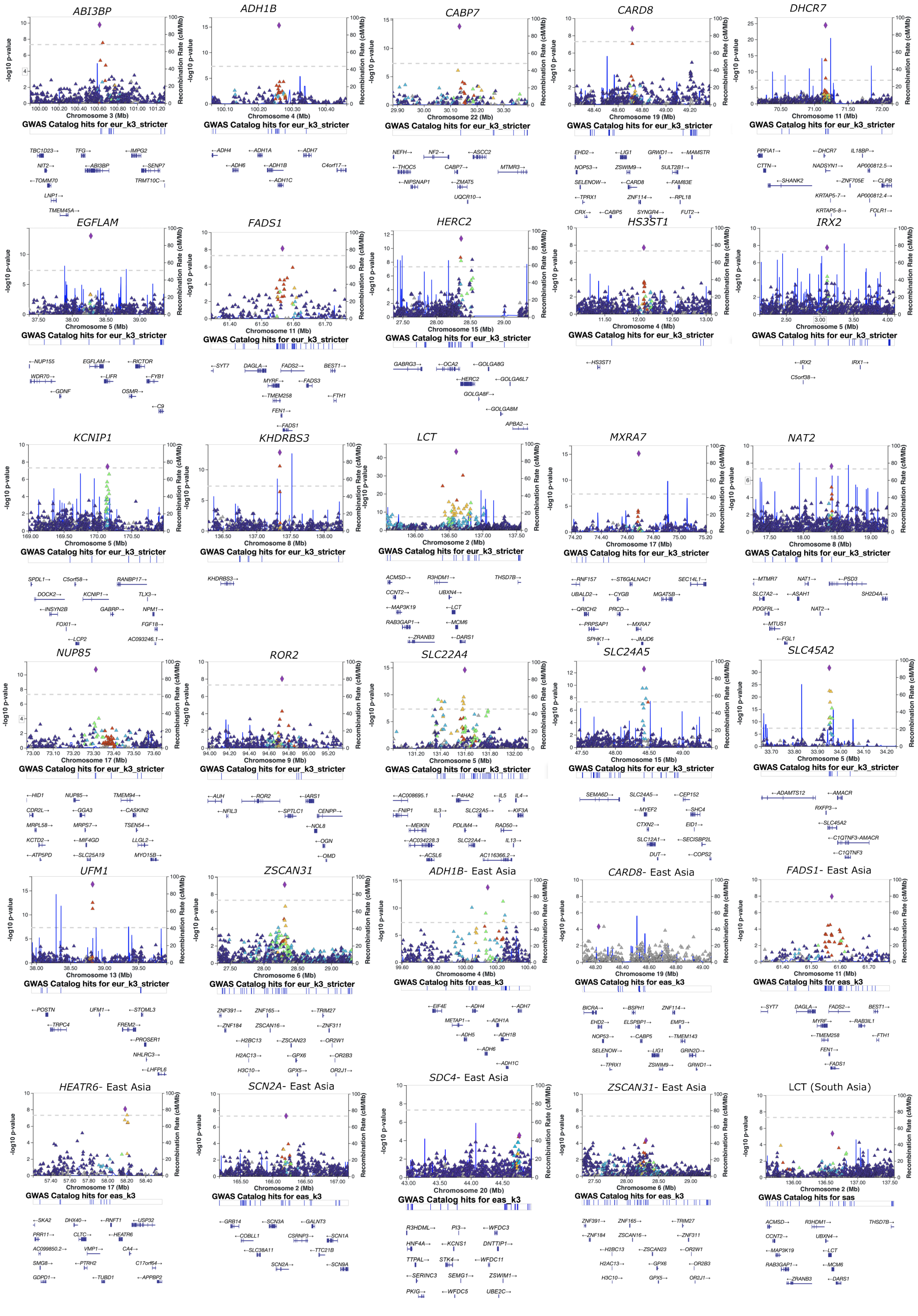
ABI3BP
ADH1B
CABP7
CARD8
DHCR7
EGFLAM
FADS1
HERC2
HS3ST1
IRX2
KCNIP1
KHDRBS3
LCT
MXRA7
NAT2
NUP85
ROR2
SLC22A4
SLC24A5
SLC45A2
UFM1
ZSCAN31
ADH1B- East Asia
CARD8- East Asia
FADS1- East Asia
HEATR6- East Asia
SCN2A- East Asia
SDC4- East Asia
ZSCAN31- East Asia
LCT (South Asia)

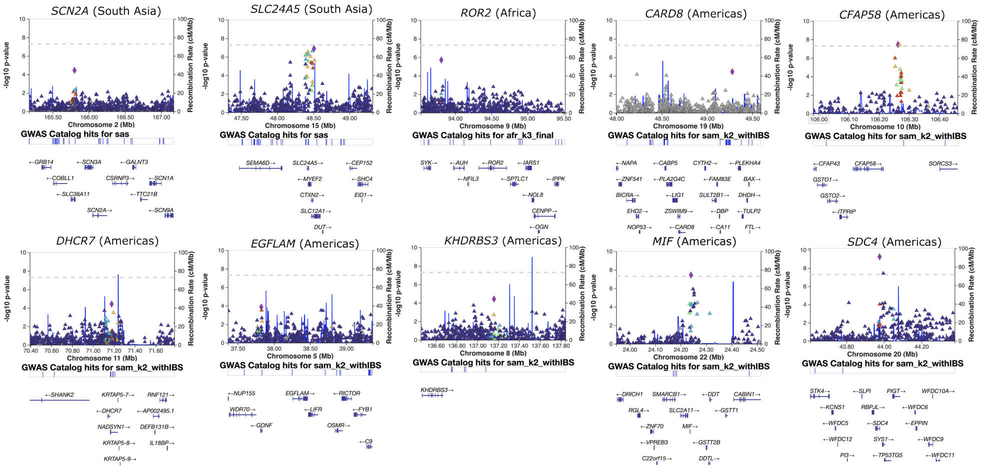


Supplementary Figure S8: LocusZoom plots for significant and replicating peaks in all 5 geographic regions

. Points are colours by LD in the relevant continental population with respect the lead SNP of each peak. Loci are labelled with the nearest gene in most cases, or the putative causal one if known.

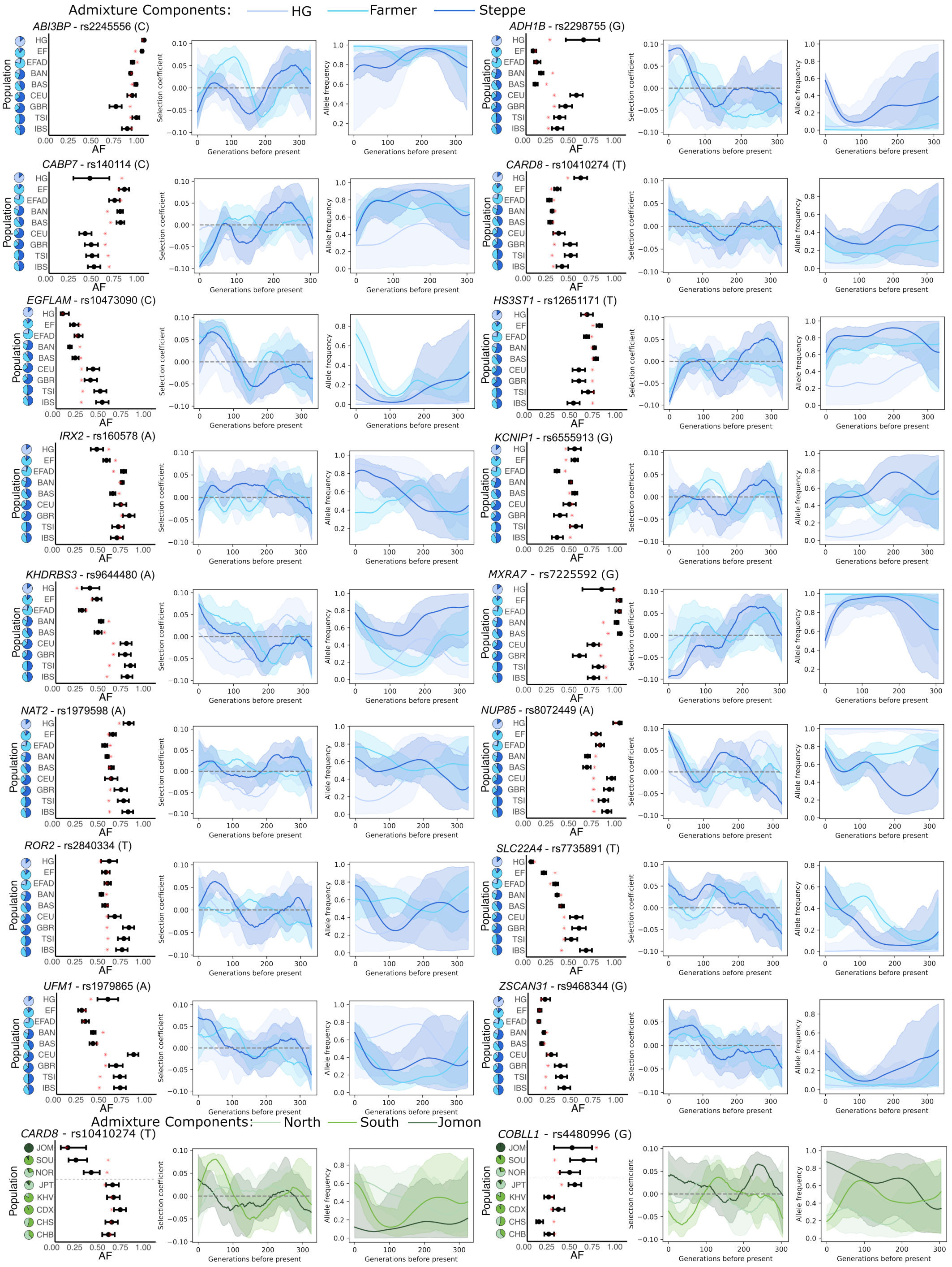
Admixture Components: HG Farmer Steppe
ABI3BP - rs2245556 (C)
ADH1B - rs2298755 (G)
CABP7 - rs140114 (C)
CARD8 - rs10410274 (T)
EGFLAM - rs10473090 (C)
HS3ST1 - rs12651171 (T)
IRX2 - rs160578 (A)
KCNIP1 - rs6555913 (G)
KHDRBS3 - rs9644480 (A)
MXRA7 - rs7225592 (G)
NAT2 - rs1979598 (A)
NUP85 - rs8072449 (A)
ROR2 - rs2840334 (T)
SLC22A4 - rs7735891 (T)
UFM1 - rs1979865 (A)
ZSCAN31 - rs9468344 (G)
Admixture Components: North South Jomon
CARD8 - rs10410274 (T)
COBLL1 - rs4480996 (G)
Population
AF
Selection coefficient
Allele frequency
Generations before present
HG
EF
EFAD
BAN
BAS
CEU
GBR
TSI
IBS
JOM
SOU
NOR
JPT
KHV
CDX
CHS
CHB

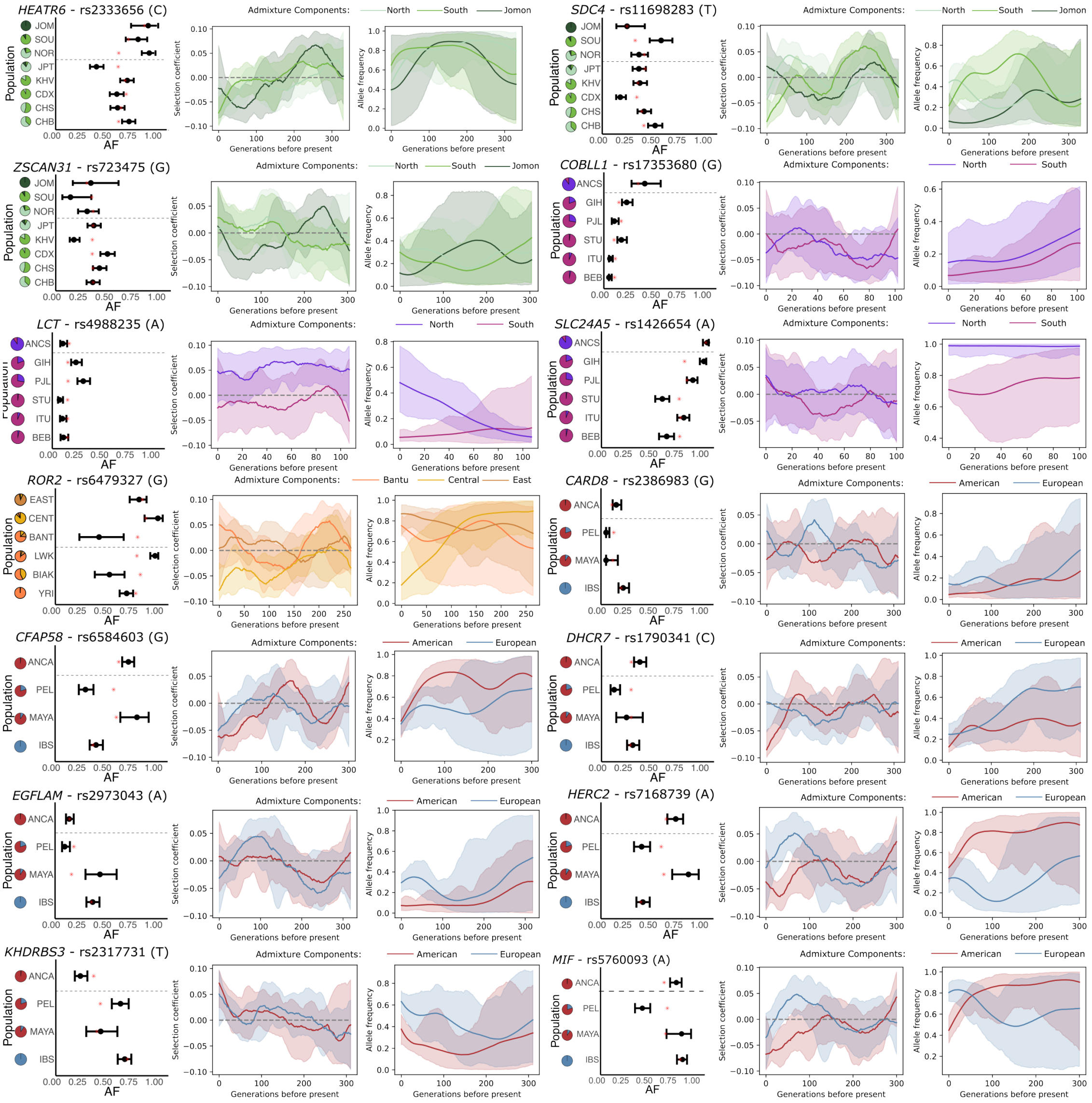


Supplementary Figure S9: Observed AF, Selection coefficient, and AF Trajectory plots for Bonferroni-significant and replication signals not highlighted in the main text.
Within each region, loci are ordered alphabetically by causal/nearest gene. We plotted the information for the derived allele of the lead SNP in each peak. In the frequency plots, the error bars represent the 95% confidence intervals calculated using the Agresti-Coull method, while the red asterisk indicates the fitted frequency from the admixture model and the dotted line separates ancient (top) from present-day (bottom) groups. The pie charts indicate the average proportion of each admixture component for each population.

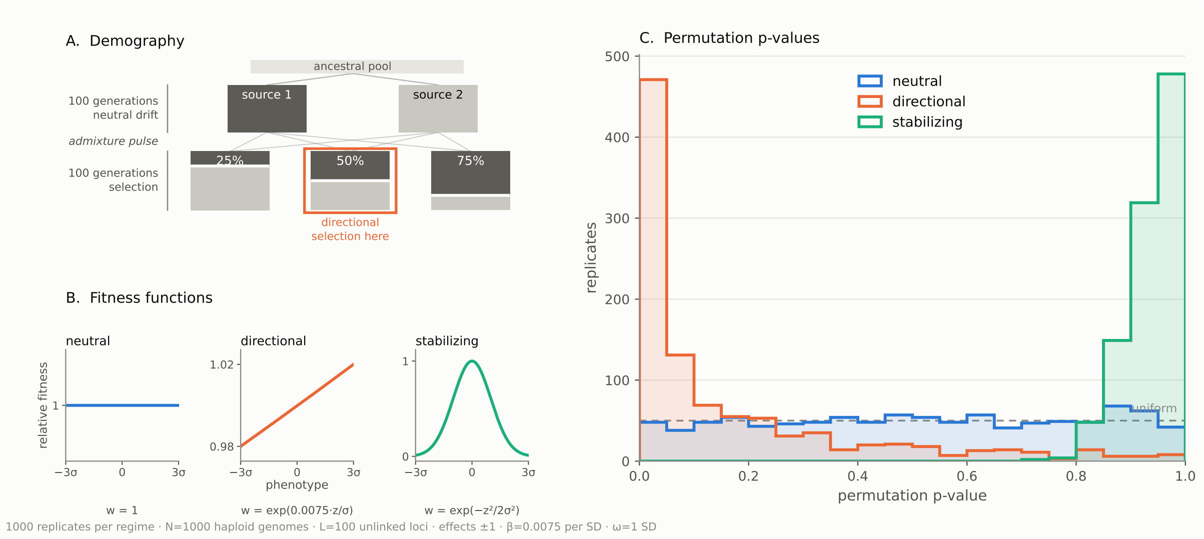


Supplementary Figure S10: Simulated polygenic selection. A) We simulate a simple demographic model of admixture with B) Three different selective regimes in the admixed populations. C) Histogram of P-values from the permutation test in the three scenarios.

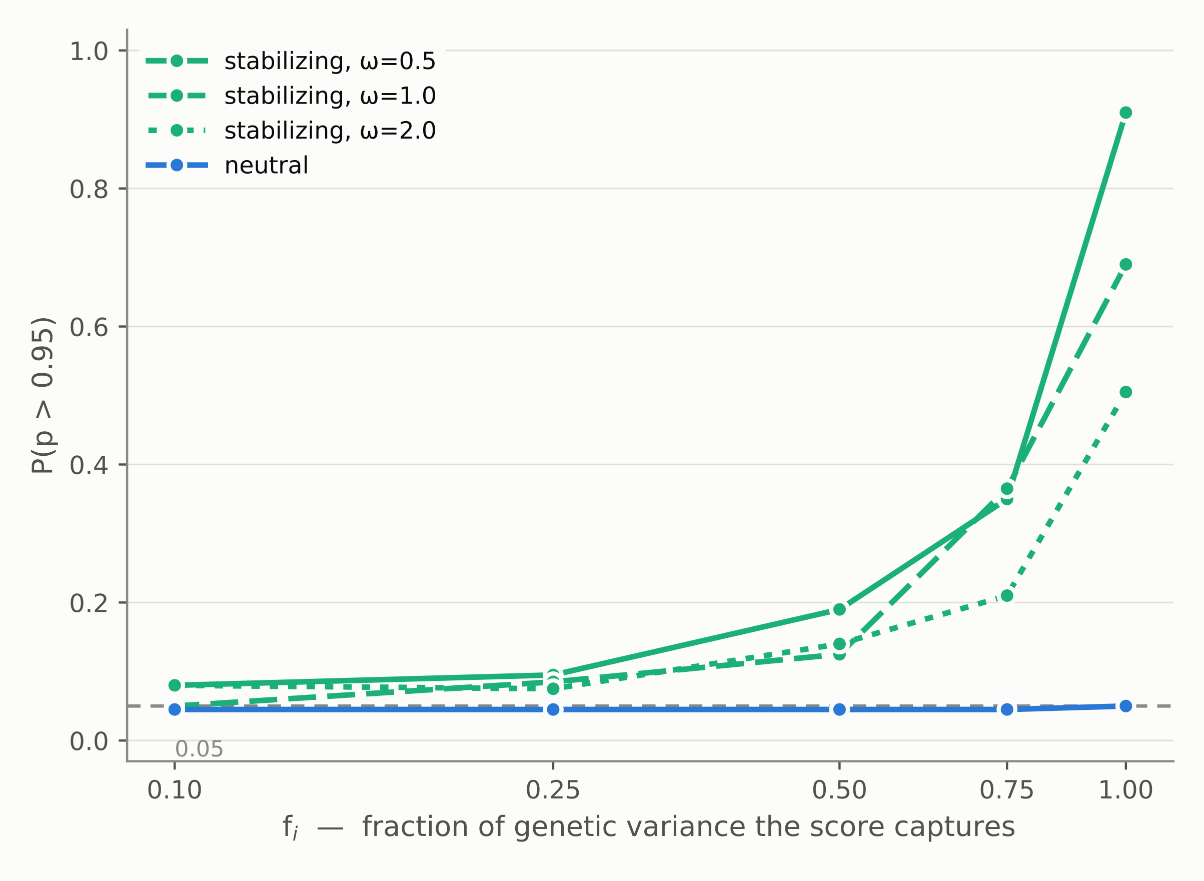


Supplementary Figure S11: Power to detect stabilizing selection. We simulated a scenario designed to match our East Asian Test population: 3 sources and 8 populations matching the real admixture proportions, N=100 chromosomes sampled in each population, 1000 trait-associated loci, 200 generations since admixture, $F_{ST} = 0.05$ between source populations. The x-axis shows how much of the underlying genetic variance is captured by the 1000 sampled loci, and the y-axis shows power to detect stabilizing selection for different fitness function widths at $\alpha = 0.05$.

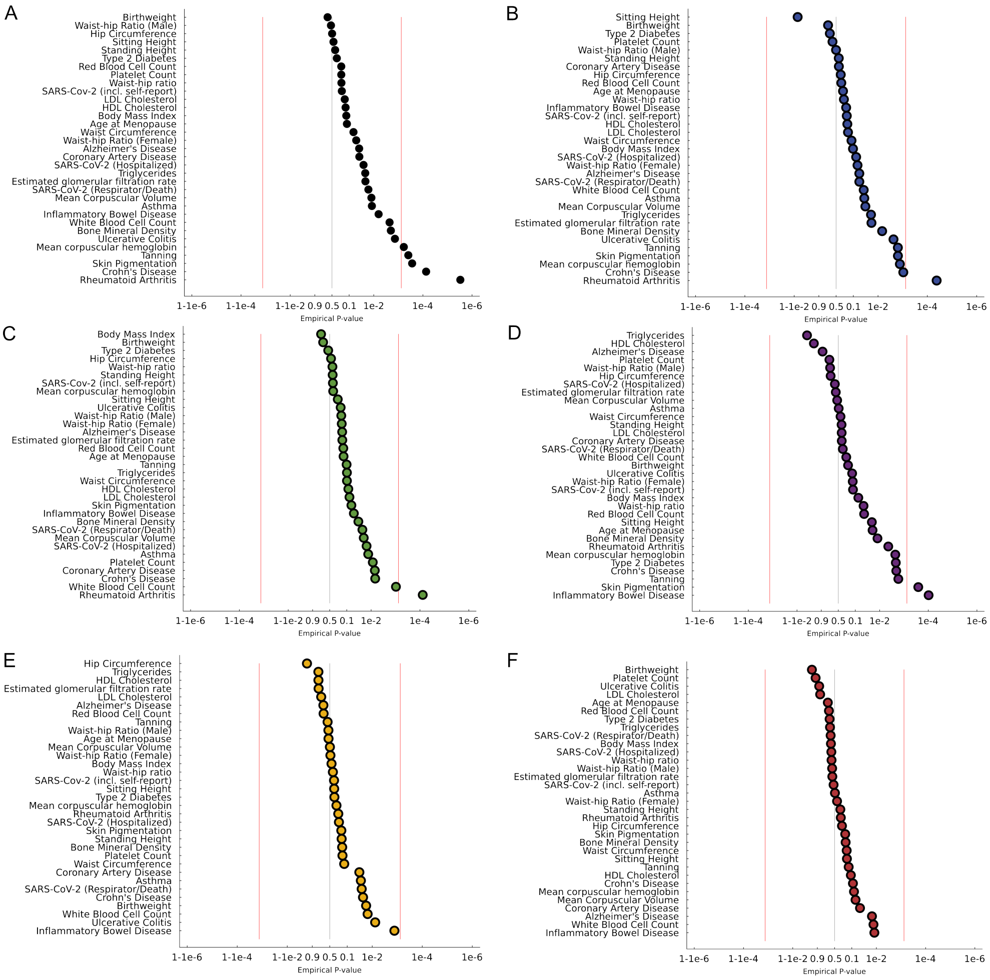


Supplementary Figure S12: Results of the polygenic tests run using European GWAS.
A) Joint results for all 5 regions, and region-specific results for all 33 traits in B) Europe, C) East Asia, D) South Asia, E) Africa, and F) the Americas.

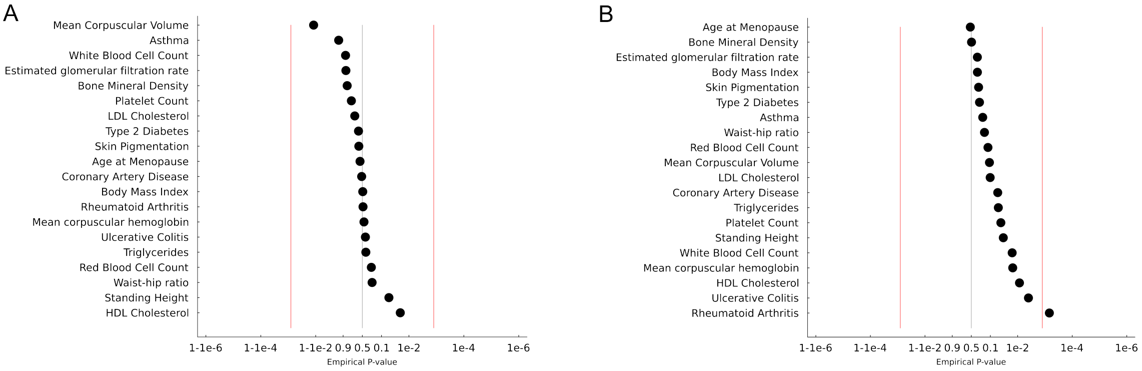


Supplementary Figure S13: Results of the joint polygenic test run using East Asian GWAS.
A) A joint analysis without the East Asia region, which might be influenced by population stratification in the underlying GWAS, B) and the full joint analysis.

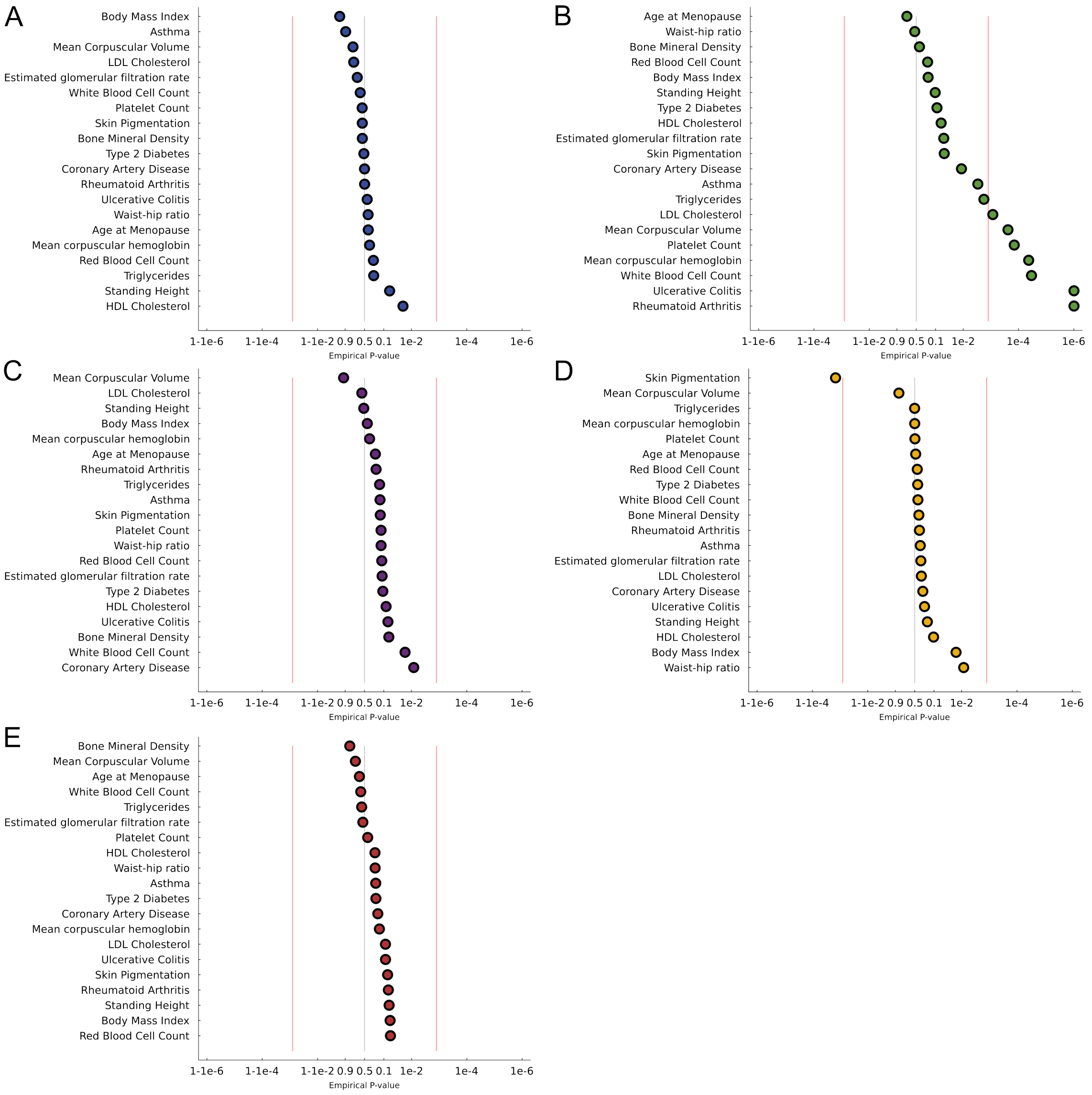


Supplementary Figure S14: Results of the polygenic tests run using East Asian GWAS.
Region-specific results for all 20 traits in A) Europe, B) East Asia, C) South Asia, D) Africa, and E) the Americas.

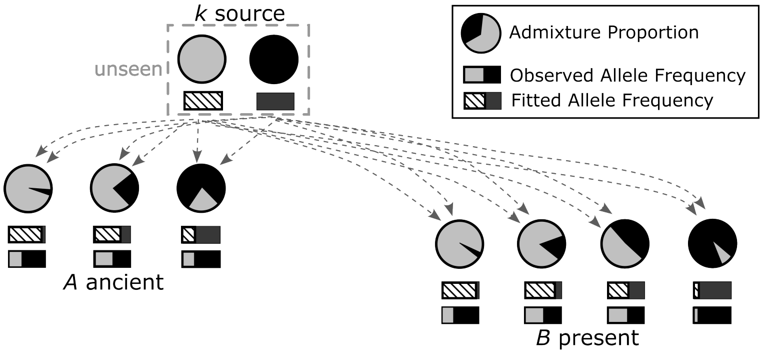


Supplementary Figure S15: We developed a test for selection by modeling the demographic history of populations.
We fit expected allele frequencies based on proportions of admixture of source populations in both ancient and modern populations (Methods).

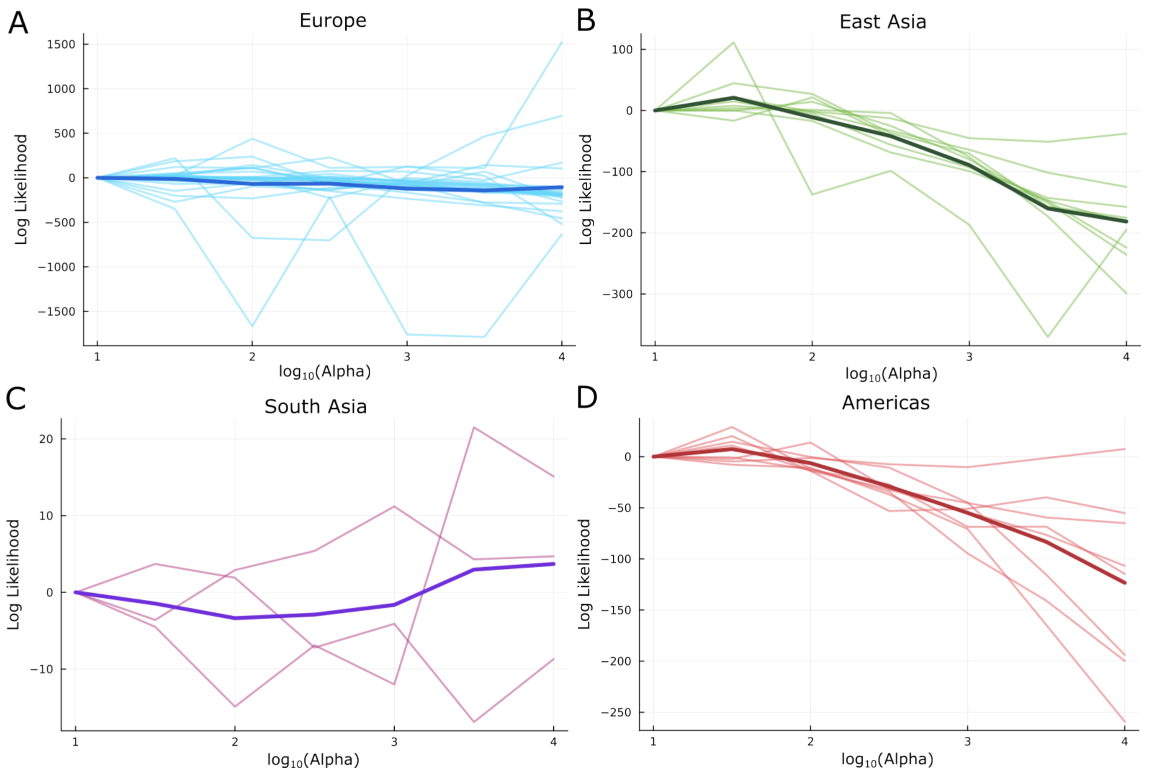


Supplementary Figure S16: Log Likelihoods for the time series selection scan plotted against a range of smoothing parameters for A) Europe, B) East Asia, C) South Asia, and D) the Americas. We did not include Africa for this as there was only one variant we were testing there. Each lighter line is a variant, while the dark one is the average of all variants. For each line the likelihood is transformed by subtracting the likelihood at $\alpha = 1$, such that each starts at zero. Based on this, we used $\log_{10} \alpha = 1.5$ for all analyses in the paper.

### Supplementary Tables

| Locus | Variant(s) | Novelty |
|---|---|---|
| *SLC24A5* | Europe (South Asia) | Known |
| *LCT* | Europe (South Asia) | Known |
| *HERC2* | Europe (Americas) | Known |
| *SLC45A2* | Europe | Known |
| *SLC22A4* | Europe | Known |
| *DHCR7* | Europe (Americas) | Known |
| *ADH1B* | East Asia + Europe | Known |
| *FADS1* | East Asia + Europe | Known |
| *HEATR6* | East Asia | Novel |
| *COBLL1* | East Asia (South Asia) | Novel |
| *IRX2* | Europe | Novel |
| *KCNIP1* | Europe | Novel |
| *ABI3BP* | Europe | Novel |
| *MXRA7* | Europe | Novel |
| *CARD8* | Europe (East Asia, Americas) | Novel |
| *EGFLAM* | Europe (Americas) | Novel |
| *HS3ST1* | Europe | Novel |
| *CABP7* | Europe | Novel |
| *NAT2* | Europe | Novel |
| *UFM1* | Europe | Novel |
| *ROR2* | Europe (Africa) | Novel |
| *KHDRBS3* | Europe (Americas) | Novel |
| *NUP85* | Europe | Novel |
| *ZSCAN31* | Europe (East Asia) | Novel |
| *SDC4* | Americas (East Asia) | Novel |
| *MIF* | Americas | Novel |
| *CFAP58* | Americas | Novel |
| *HLA-DQB1* | Europe (East Asia) | Known |
| *HLA-B* | Europe + Americas | Known |
| *HLA-F* | Americas (East Asia) | Known |
| *NOTCH4* | Europe | Known |

Supplementary Table S1: Table of all loci passing a Bonferroni multiple testing correction in at least one region. Regions in parentheses indicate those that at least nominally replicate the peak (Methods). Novelty indicates whether this locus has been previously identified in a selection scan.

| Trait | SNPs | AFR. P | EAS P | EUR P | SAS P | AMR P | Joint | NonEas |
|---|---|---|---|---|---|---|---|---|
| Asthma | 64 | 0.31 | 0.003 | 0.90 | 0.20 | 0.14 | 0.19 | 0.93 |
| Bone Mineral Density | 190 | 0.35 | 0.38 | 0.58 | 0.85 | 0.066 | 0.49 | 0.86 |
| Body Mass Index | 264 | 0.016 | 0.19 | 0.94 | 0.060 | 0.40 | 0.30 | 0.48 |
| Coronary Artery Disease | 79 | 0.25 | 0.012 | 0.50 | 0.17 | 0.0083 | 0.054 | 0.53 |
| Est. glomerular filtration rate | 61 | 0.30 | 0.05 | 0.73 | 0.57 | 0.12 | 0.30 | 0.87 |
| HDL Cholesterol | 401 | 0.10 | 0.063 | 0.020 | 0.21 | 0.083 | $8.6 \times 10^{-3}$ | 0.021 |
| Standing Height | 1762 | 0.17 | 0.10 | 0.062 | 0.065 | 0.53 | 0.034 | 0.054 |
| LDL Cholesterol | 212 | 0.29 | $8.5 \times 10^{-4}$ | 0.80 | 0.088 | 0.60 | 0.10 | 0.73 |
| Triglycerides | 343 | 0.50 | $1.8 \times 10^{-3}$ | 0.23 | 0.61 | 0.14 | 0.051 | 0.37 |
| Mean corpuscular hemoglobin | 324 | 0.50 | $4.3 \times 10^{-5}$ | 0.33 | 0.15 | 0.33 | 0.015 | 0.44 |
| Mean corpuscular volume | 330 | 0.87 | $2.4 \times 10^{-4}$ | 0.81 | 0.77 | 0.91 | 0.11 | 0.99 |
| Age at Menopause† | 45 | 0.46 | 0.77 | 0.36 | 0.67 | 0.20 | 0.54 | 0.58 |
| Platelet Counts | 875 | 0.49 | $1.4 \times 10^{-4}$ | 0.59 | 0.38 | 0.13 | 0.041 | 0.80 |
| Rheumatoid Arthritis | 248 | 0.34 | $< 10^{-6}$* | 0.49 | 0.069 | 0.19 | $6.9 \times 10^{-4}$ | 0.47 |
| Red Blood Cell Count | 453 | 0.40 | 0.19 | 0.24 | 0.058 | 0.12 | 0.12 | 0.23 |
| Skin Pigmentation† | 31 | 0.9993 | 0.049 | 0.58 | 0.073 | 0.13 | 0.27 | 0.63 |
| Type II Diabetes | 353 | 0.39 | 0.089 | 0.53 | 0.19 | 0.11 | 0.25 | 0.64 |
| Ulcerative Colitis† | 59 | 0.22 | $< 10^{-6}$* | 0.40 | 0.087 | 0.071 | 0.004 | 0.39 |
| White Blood Cell Counts | 360 | 0.38 | $3.4 \times 10^{-5}$ | 0.65 | 0.64 | 0.017 | 0.016 | 0.88 |
| Waist-Hip Ratio† | 51 | 0.0085 | 0.56 | 0.37 | 0.21 | 0.13 | 0.16 | 0.22 |

Supplementary Table S2: Results of testing for polygenic selection on 20 traits with East Asian-ancestry GWAS. Fields include the trait abbreviation, number of variants included in the test, and the p-values in Africa, East Asia, Europe, South Asia, and the Americas, as well as those for the joint analysis over all regions, and that excluding East Asians. Loci for all traits were those that passed a $p < 1e-8$ threshold in the original GWAS except for † traits, where we used $1e-6$ instead. *passes Bonferroni correction < 0.05.

| Trait | SNPs | AFR. P | EAS P | EUR P | SAS P | AMR P | Joint | NonEur |
|---|---|---|---|---|---|---|---|---|
| Alzheimer's Disease | 42 | 0.72 | 0.15 | 0.057 | 0.015 | 0.89 | 0.039 | 0.190 |
| Asthma | 586 | 0.028 | 0.013 | 0.036 | 0.48 | 0.48 | 0.012 | 0.020 |
| Bone Mineral Density | 1080 | 0.16 | 0.033 | $6.8 \times 10^{-3}$ | 0.18 | 0.012 | $2 \times 10^{-3}$ | 0.060 |
| Body Mass Index | 831 | 0.42 | 0.78 | 0.10 | 0.64 | 0.075 | 0.13 | 0.97 |
| Birthweight | 115 | 0.018 | 0.73 | 0.76 | 0.94 | 0.20 | 0.67 | 0.75 |
| Crohn's Disease | 263 | 0.023 | $6.8 \times 10^{-3}$ | $9.4 \times 10^{-4}$ | 0.090 | $2.1 \times 10^{-3}$ | $7.4 \times 10^{-5}$ | $4.4 \times 10^{-3}$ |
| Covid-19 (Respirator/Death) | 69 | 0.026 | 0.023 | 0.057 | 0.65 | 0.33 | 0.026 | 0.053 |
| Covid-19 (Hospitalized) | 70 | 0.21 | 0.015 | 0.077 | 0.63 | 0.64 | 0.20 | 0.69 |
| Covid-19 (incl. self-report) | 43 | 0.34 | 0.37 | 0.18 | 0.54 | 0.12 | 0.017 | 0.038 |
| Coronary Artery Disease | 391 | 0.033 | 0.007 | 0.39 | 0.046 | 0.36 | 0.038 | 0.003 |
| Est. glomerular filtration rate | 281 | 0.821 | 0.15 | 0.018 | 0.59 | 0.61 | 0.022 | 0.63 |
| HDL Cholesterol | 1238 | 0.82 | 0.086 | 0.18 | 0.11 | 0.95 | 0.14 | 0.20 |
| Standing Height | 4102 | 0.17 | 0.38 | 0.39 | 0.29 | 0.37 | 0.37 | 0.51 |
| Hip Circumference | 810 | 0.94 | 0.44 | 0.32 | 0.25 | 0.76 | 0.49 | 0.82 |
| Inflammatory Bowel Disease | 310 | $1.3 \times 10^{-3}$ | 0.051 | 0.20 | 0.012 | $9.7 \times 10^{-5}$ | $6.3 \times 10^{-3}$ | $2.8 \times 10^{-3}$ |
| LDL Cholesterol | 1493 | 0.77 | 0.077 | 0.17 | 0.87 | 0.36 | 0.15 | 0.44 |
| Triglycerides | 1371 | 0.82 | 0.098 | 0.019 | 0.67 | 0.97 | 0.022 | 0.48 |
| Mean corpuscular hemoglobin | 478 | 0.27 | 0.36 | $1.3 \times 10^{-3}$ | 0.081 | $2.3 \times 10^{-3}$ | $6 \times 10^{-4}$ | 0.32 |
| Mean corpuscular volume | 1846 | 0.49 | 0.020 | 0.033 | 0.070 | 0.55 | 0.012 | 0.029 |
| Age at Menopause | 143 | 0.55 | 0.14 | 0.26 | 0.73 | 0.020 | 0.12 | 0.62 |
| Platelet Counts | 2160 | 0.16 | $8.6 \times 10^{-3}$ | 0.64 | 0.91 | 0.79 | 0.21 | 0.034 |
| Rheumatoid Arthritis | 621 | 0.23 | $7.7 \times 10^{-5}$ | $4.2 \times 10^{-5}$ | 0.279 | $4.4 \times 10^{-3}$ | $3 \times 10^{-6}$ | $1.4 \times 10^{-4}$ |
| Red Blood Cell Count | 1451 | 0.72 | 0.14 | 0.31 | 0.71 | 0.043 | 0.21 | 0.57 |
| Sitting Height | 2640 | 0.33 | 0.23 | 0.99 | 0.16 | 0.021 | 0.43 | 0.34 |
| Skin Pigmentation | 657 | 0.17 | 0.064 | $1.6 \times 10^{-3}$ | 0.19 | $2.6 \times 10^{-4}$ | $2.7 \times 10^{-4}$ | 0.13 |
| Type II Diabetes | 423 | 0.33 | 0.56 | 0.72 | 0.69 | $2.2 \times 10^{-3}$ | 0.32 | 0.89 |
| Tanning | 654 | 0.59 | 0.10 | $1.6 \times 10^{-3}$ | 0.13 | $1.7 \times 10^{-3}$ | $3.9 \times 10^{-4}$ | 0.21 |
| Ulcerative Colitis | 202 | 0.008 | 0.18 | $2.4 \times 10^{-3}$ | 0.88 | 0.14 | $1.4 \times 10^{-3}$ | 0.22 |
| Waist Circumference | 497 | 0.13 | 0.098 | 0.12 | 0.16 | 0.41 | 0.067 | 0.14 |
| White Blood Cell Counts | 1372 | 0.015 | $9.7 \times 10^{-4}$ | 0.038 | 0.013 | 0.24 | $2.3 \times 10^{-3}$ | $1.5 \times 10^{-4}$ |
| Waist-Hip Ratio | 1362 | 0.37 | 0.39 | 0.24 | 0.62 | 0.046 | 0.21 | 0.78 |
| Waist-Hip Ratio (Females) | 978 | 0.47 | 0.16 | 0.068 | 0.40 | 0.13 | 0.052 | 0.49 |
| Waist-Hip Ratio (Males) | 338 | 0.56 | 0.17 | 0.50 | 0.60 | 0.77 | 0.55 | 0.56 |

Supplementary Table S3: Results of testing for polygenic selection on 33 traits with European-ancestry GWAS. Fields include the trait, number of variants included in the test, and the p-values in Africa, East Asia, South Asia, Europe, and the Americas, as well as the joint analysis. Loci for all traits were those that passed a $p < 1e - 8$ threshold in the original GWAS. *passes two-sided Bonferroni correction $<$ 0.05.

[11] Jessie Kulsuptrakul, Elizabeth A Turcotte, Michael Emerman, and Patrick S Mitchell. A human-specific motif facilitates CARD8 inflammasome activation after HIV-1 infection. *eLife*, 12, 2023.

[12] Tugay Karakaya, Marta Slaufova, Michela Di Filippo, et al. CARD8: A Novel Inflammasome Sensor with Well-Known Anti-Inflammatory and Anti-Apoptotic Activity. *Cells*, 13(12), 2024. doi: 10.3390/cells13121032.

[13] Zheng-Zheng Li, Wen-Long Zhao, Gui-Shuan Wang, et al. The novel testicular enrichment protein Cfap58 is required for Notch-associated ciliogenesis. *Bioscience reports*, 40(1), 2020.

[14] Yibin Xu, Armand 3rd Bankhead, Xiaoli Tian, et al. Deletion of Glutathione S-Transferase Omega 1 Activates Type I Interferon Genes and Downregulates Tissue Factor. *Cancer research*, 80(17): 3692–3705, 2020.

[15] D M Escobar-García, L M Del Razo, L C Sanchez-Peña, et al. Association of glutathione S-transferase Ω 1-1 polymorphisms (A140D and E208K) with the expression of interleukin-8 (IL-8), transforming growth factor beta (TGF-$\beta$), and apoptotic protease-activating factor 1 (Apaf-1) in

humans chronically exposed to arsenic in drinking water. *Archives of Toxicology*, 86(6):857–868, 2012.

[35] Lin Poyraz, Laura L Colbran, and Iain Mathieson. Predicting functional consequences of recent natural selection in britain. *Molecular Biology and Evolution*, 41:msae053, 2024.
